# Genome duplication and transposon mediated gene alteration shapes the pathogenicity of *Rhizoctonia solani* AG1-IA

**DOI:** 10.1101/2022.07.01.498367

**Authors:** Aleena Francis, Srayan Ghosh, Kriti Tyagi, V. Prakasam, Mamta Rani, Nagendra Pratap Singh, Amrita Pradhan, R. M. Sundaram, C. Priyanka, G.S. Laha, C. Kannan, M.S. Prasad, Debasis Chattopadhyay, Gopaljee Jha

## Abstract

*Rhizoctonia solani* AG1-IA is a polyphagous basidiomycete fungal pathogen that causes sheath blight disease in rice. In a high-quality genome assembly-based analysis, we report a recent whole genome duplication in *R. solani* AG1-IA. Duplicated syntenic gene blocks showed presence of district clusters of transposable elements (TEs), which introduced disruption in the continuity of synteny and caused alterations in gene structures. Genome duplication followed by TE-mediated gene structure alterations caused neofunctionalization of genes associated with pathogenicity, as experimentally shown by variation in expression patterns and their involvement during plant colonization. High throughput genome sequencing of forty-two rice field isolates of *R. solani* AG1-IA from different agro-climatic zones of India profiled the population genetic structure of the Indian isolates and classified those into three distinct groups and a subgroup of admixture, emphasizing exchange of genetic material under field conditions. Genetic diversity analysis of this population predicted the regions that are that are targets for diversifying and purifying selections. Experimental evidence showed that the genes undergoing diversifying and purifying selections were essential for pathogenicity. Together, our data and the analysis revealed profound impact of genome duplication and the transposable elements on genomic diversity and evolution that shaped the pathogenicity of *R. solani* AG1- IA.

## Introduction

*Rhizoctonia solani,* a basidiomycetes necrotrophic fungal pathogen, infects a broad range of plant species, including several economically important crops, such as rice, tomato, potato, maize, barley, turf grass etc^1–3^. The polyphagous nature enables it to survive several years in the soil, even in absence of primary host. *R. solani* strains have been classified into 14 different anastomosis group (AG) i.e. AG1 (which is further divided into intraspecific groups: IA, IB, IC, ID, IF and IE) to AG13 and AGBI^1, 4^. Although classified into the same taxonomic group, strains belonging to different AGs are mostly sexually incompatible. *R. solani* strains exhibit large morphological and pathological diversity^5, 6^ and they also differ in karyotype banding pattern, number of nuclei, as well as chromosome number per somatic cells^1, 7^. Therefore, these features emphasize the need to understand the genomic diversity and evolutionary relationship between *R. solani* strains belonging to different AGs.

The genomes of AG1-IA^8, 9^ , AG1-IB^10^, AG2-2IIIB^11^, AG3^12^ and AG8^13^ strains of *R. solani* have been sequenced These studies have catalogued various pathogenicity-associated genes, PHIbase homologs, effectors, cell wall degrading enzymes (CAZymes) and secondary metabolites encoded in different *R. solani* genomes. Also, the core genes that are conserved in different AGs and unique genes present in particular AG strain have been identified^11, 13^. Recent analyses have suggested the AG1-IA strains to be diverse from other AGs^8, 14^ and they are enriched in homogalacturonan/pectin modification genes and have expanded pathogenicity-associated gene families^7, 8^.

Generally, high degree of heterozygosity (due to their coenocytic nature), nucleotide variations (SNPs, Indels), large scale chromosomal rearrangements (insertion, inversion, and deletion, leading to structural and organizational changes in the chromosome) and presence of accessory/mobile chromosomes are observed in different strains of pathogenic fungi ^15–17^. They are enriched in repetitive DNAs^16^ which serve as a cradle for the evolution of pathogenicity-associated traits^18^. The transposons and other mobile elements play important role in the evolution of the fungal pathogens and modulate host specificity as well as aggressiveness^19, 20^. However, due to use of short read-based sequencing technology and assembly of the genome into numerous fragmented contigs, repetitive DNA content, genome complexity and evolutionary relationship amongst different AG strains of *R. solani* remain elusive. In this study, we have produced a very high quality genome assembly using long read technology of a pathogenic strain (BRS1, a rice isolate from India) of *R. solani* AG1-IA that causes sheath blight disease in rice^21^. The 44.5 MB genome has been assembled into 74 contigs and a database is created to facilitate the interactive analyses of the genome. Computational analysis has unravelled the evolutionary relationship and divergence of different AG strains of *R. solani*, in evolutionary time scale. Moreover, relatively large transposon repertoire and segmental gene duplication events leading to neo-functionalization of genes have been identified during the current study. Further, through genome sequencing and comparative sequence analysis of different *R. solani* AG1-IA isolates (n=42; collected from rice-growing regions across India) we have identified genes undergoing positive and purifying selection in *R. solani*. Using some of the selected genes undergoing positive as well as purifying selction, the importance of neofunctionalization in pathogenicity of *R*. *solani* have been established.

## Results

### Assembly and annotation of *R. solani* AG1-IA (strain BRS1) genome

We adopted SMRT long-read sequencing platform to sequence and assemble the genome of *R. solani* AG1-IA strain BRS1^22^ (pathogenic Indian isolate). 13.73 Gb sequence data generated using PacBio Sequel II platform with an average read length of 10.3 Kb was assembled by the diploid-aware assembler FALCON to generate primary and associated contigs and followed by diploid assembler FALCON-Unzip to generate 74 primary contigs. These primary contigs were sequence-corrected by using about 6 Gb Illumina whole-genome sequencing (WGS) short-read data following Pilon to finally produce a 44,527,001 bp genome assembly in 74 contigs with an average length of 601,716 bp and N50 length of 2,014,351 bp (Table S1). Notably, more than half of the genome assembly was covered in nine fragments and compared to the previous AG1-IA assembly^9^ which showed 0.12% heterozygosity, six-times higher heterozygosity (0.76%) was observed in the BRS1 genome assembly. The heterozygous bases are evenly distributed throughout the genome, irrespective of high- or low-density gene regions (Fig. 1). Annotation of repeat sequences predicted a presence of a total 23.35% interspersed repeat sequence with a huge proportion of retroelement (95% being from Gypsy family) spreading over 10.75% of the assembly (Table S2). Most of the contigs showed the presence of transposon elements (TEs) being interspersed with the protein-coding genes (Fig. 1).

**Fig. 1.**
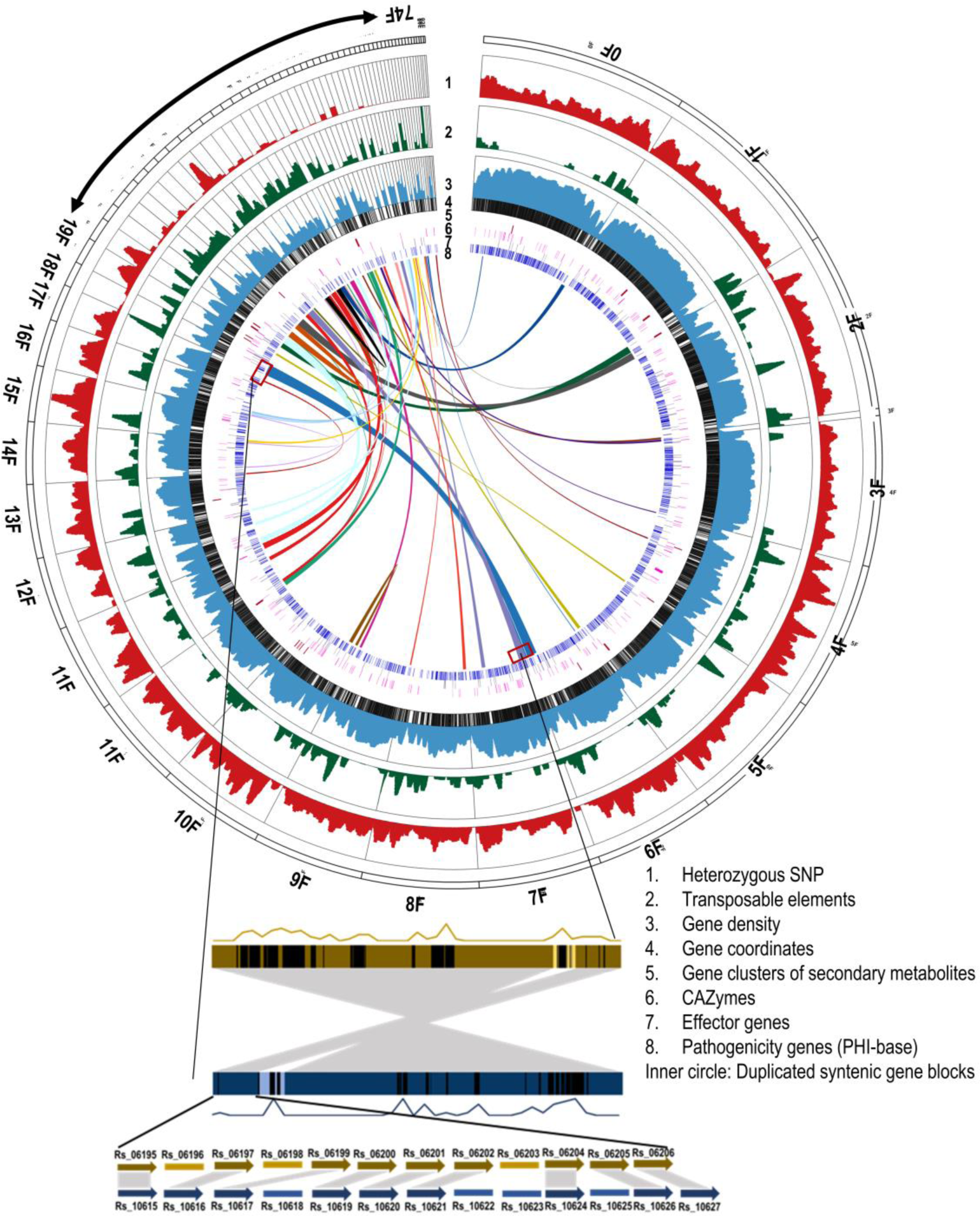
Genomic features of the *R. solani* AG1-IA genome assembly. The Circose diagram showing the density of heterozygous SNPs, genes and transposons in 100 kb with 10 kb sliding windows were presented. The scale used for heterozygous SNPs was 0-3000, for the percentage of transposable elements 0-100, and for the percentage of protein-coding genes was 0-100. Positions of genes, secondary metabolite-associated gene clusters, CAZymes, effectors genes and PHI-base genes were marked by coloured lines according to their coordinates. Each duplicated syntenic gene blocks are denoted with different colours. Contigs names are displayed around the circle. The synteny between two duplicated blocks marked with red boxes are shown below. Density and the coordinates of TEs in each block are shown by the density curve and black lines. Light colours within the duplicated blocks denote non-syntenic regions.

Repeat-masked assembly was annotated for protein-coding genes following *ab initio*, homology-based, and transcriptome-based approaches and finally combining them using Evidence modeller predicted 11,902 high-confidence genes and 14,261 open reading frames (ORFs) (Table S3) belonging to 7118 unigene and 2448 multigene families with 2 to 10 members per family. Gene ontology (GO) search and manual curation by homology search in different gene family-specific databases predicted genes encoding pathogen-host interacting factors (PHI-base) (n=2,500), transcription factors (n=289), carbohydrate-active enzymes (CAZymes) (n=301), secreted proteins (759), effector proteins (n=202), proteins involved in secondary metabolism (n=177) and GPI-anchored proteins (n=56) (Fig. S3b, Table S4). Notably, genes encoding PHI-base, CAZymes, and effectors were evenly distributed throughout the genome; however, genes involved in secondary metabolism were clustered in fifteen pockets mostly located in TE-dense regions (Fig. 1). Moreover, the predicted numbers of genes encoding CAZymes and effectors in this assembly are much higher than those predicted by other assemblies of *R. solani.* Whereas, the predicted number of PHI-base orthologs is similar to that in AG2-2IIIB^11^.

The present *R. solani* AG1-IA genome assembly and protein-coding gene annotation are hosted in a web-based user-interactive Rice sheath blight (RSB) database (http://223.31.159.7/RSB/public/) along with an embedded jBrowse genome browser for visualizing the genome, and blast search options for similarity search for protein-coding genes and comparison of proteins among the different anastomosis groups of *R. solani* (Fig. S1).

### Comparative genomic analysis and evolutionary divergence of AGs in *R. solani*

Synteny analysis using the coordinates of the orthologous protein-coding genes of AG1-IA strain BRS1 and other sequenced *R. solani* AGs showed a high degree of collinearity between AG1-IA and AG8 genome assemblies with 5865 syntenic genes (Fig. S2a). AG1-IA genome showed more degree of collinearity to AG3 (7513 syntenic genes) (Fig. S2b) than to AG2- 2IIIB (6156 syntenic genes) (Fig. S2c). To delineate unique and shared protein-coding genes among the sequenced AGs, all the predicted proteomes belonging to the respective AG strains present in NCBI database were considered. Our analysis predicted 4,338 orthogroups (5,305 ORFs) out of total 20,840 orthogroups to be unique in AG1-IA (Table S5). All the sequenced *R. solani* AGs shared only 5,217 orthogroups comprising 35,929 ORFs indicating a wide genetic variation within the *R. solani* anastomosis groups. We also observed 313 unique multi- gene families (n=1280 genes) being present in *R. solani* AG1-IA (Fig. S3a). RNAseq data supported the expression of 45.6% of these AG1-IA BRS1 specific genes (Fig. S4, Table S6). We also analysed the expression of eight of the randomly selected AG1-IA unique genes using qRT-PCR during pathogenesis in rice and observed majority of them (*Rs_09732, Rs_08184, Rs_11108, Rs_01662,* and *Rs_03147*) to be upregulated during necrotrophic phase (3 dpi) of infection (Fig. S5).

Further, to determine the period of divergence of different AGs of *R. solani* we plotted the rate of synonymous substitution per synonymous site (Ks) between the orthogroups in the syntenic blocks between AG1-IA and the other sequenced AGs against the percentage of orthologous pairs. We considered the average rate in change of Ks value as 1.3x10^-8^ per year, as reported earlier in case of fungi. The maximum likelihood-based phylogenetic tree constructed using the single-copy orthogroups expectedly predicted AG1-IA and AG1-IB to be phylogenetically closest and they seems to have diverged around 28 million years ago (mya). The data suggested AG1-IA to have diverged from AG8 about 32 mya. On the other hand, AG3 and AG2-2IIIB had diverged from AG1-IA around 44 mya whereas AG3 and AG2-2IIIB had diverged from each other around 21 mya (Fig. 2).

**Fig. 2.**
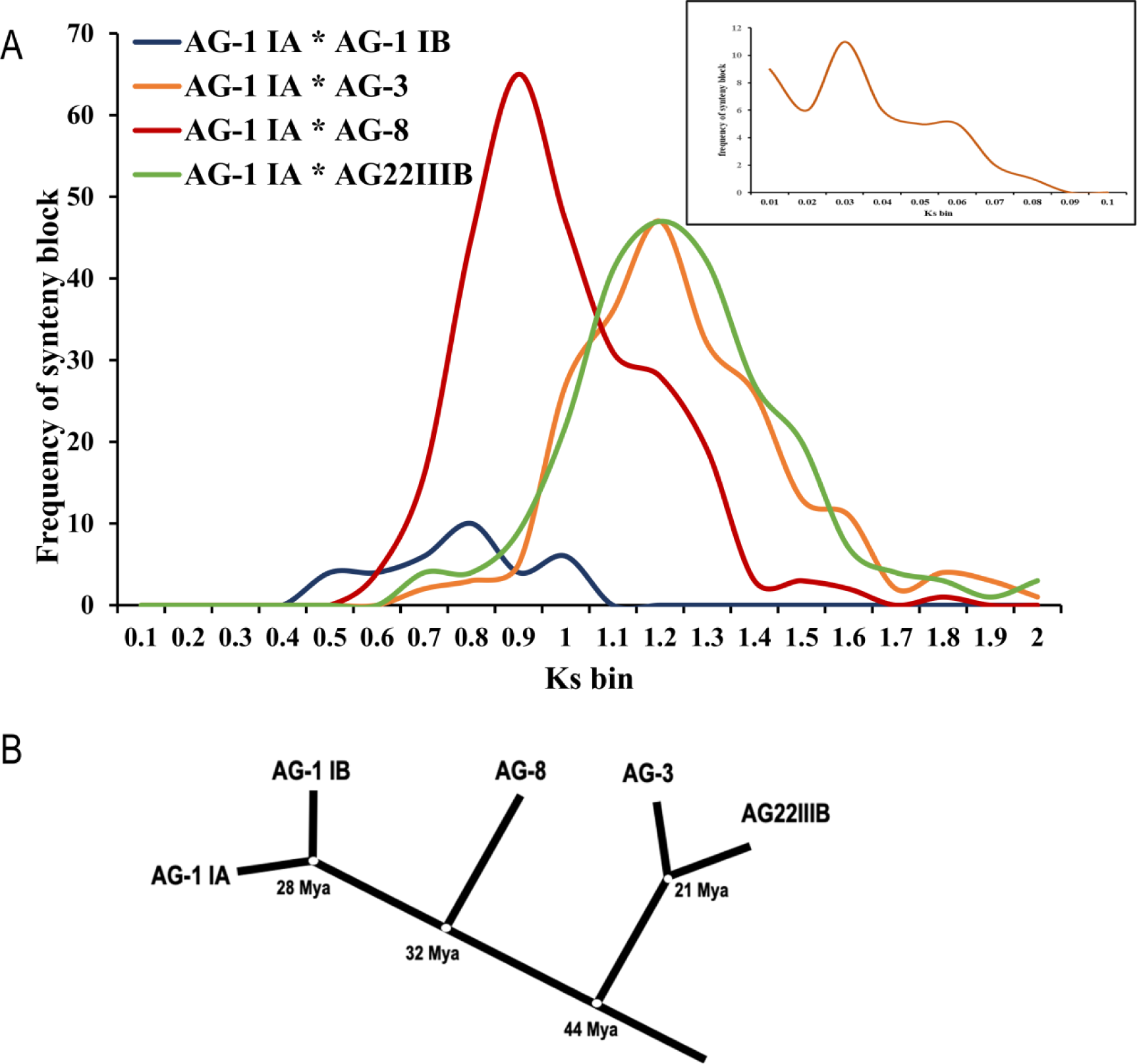
Evolutionary divergence among the AGs of *R. solani*. A. Distribution of synonymous substitution rates (Ks) of different combinations of orthologous genes of strains of *R. solani*. Ks of the orthologous gene pairs between different strains mentioned were plotted against the number of orthologous gene pairs in different colours. The inset shows distribution of Ks of the paralogous gene pairs located in the duplicated syntenic gene blocks of the genome assembly of AG1-IA. B. Phylogenetic tree of five genome-sequenced *R. solani* strains and their divergence times calculated based on Ks of the orthologous gene pairs.

### Genome duplication in *R. solani*

Genome duplication has been proposed as a strategy for adaptation and evolutionary innovation in fungus not only because it increases the gene copy number, but also it supplies new genetic material for the emergence of new functions, by mutation and selection^23, 24^. A detailed analysis revealed that about 50% of protein-coding ORFs have homologs within the *R. solani* genome and they are grouped into 2,448 (n=7143) multigene families out of which, 38.86% of the ORFs are grouped into two-member gene families. This high proportion of duplicated gene pairs prompted an investigation into whether multiple segmental duplications or an ancestral whole-genome duplication (WGD) event had occurred in *R. solani* AG1-IA. We looked for the existence of any paralogous syntenic blocks as a reminiscence genome duplication in the AG1- IA genome assembly. We identified 669 paralogous gene pairs, which can be uniquely grouped into 46 pairs of duplicated syntenic blocks having at least five and up to 58 duplicated genes in a block (Table S7). These duplicated blocks are enriched in genes encoding CAZymes, PHIbase, secreted/effectors, transcription factors, and secondary metabolite-related proteins. To rule out the possibility that the contigs hosting the syntenic blocks might had arise due to multinucleated cells, we mapped the long reads as well as the short reads of WGS and the RNAseq reads to these contigs. Almost 94% of the long reads and 90% of the short reads and 97.36% of RNASeq reads were uniquely mapped on the respective contigs (Table S8). Apart from that, interspersed genes between the paralogous blocks and the genes flanking these blocks in the corresponding contigs were also different, indicating these contigs to be unique. Together, the duplicated regions covered approximately 15% of the genome assembly and spanned over 55 out of 74 scaffolds. The duplicated genes in each of these regions are found in the same order and orientation, providing evidence of an ancestral duplicated state for these regions (Fig. 1). Alternatively, if the 46 duplicated blocks were the resultant of sequential segmental duplication, some of the early duplicated blocks would have been part of later events of segmental duplication and according to Poisson distribution would have resulted in 8 triplets within 46 duplicated blocks. We have detected 4 triplicate blocks with minimum five genes (Fig. S6) with a moderate probability (*p* = 0.057) indicating sequential segmental duplication along with whole genome duplication might have happened in *R. solani* AG1-IA. Analysis of synonymous substitution rates of the paralogous gene groups indicated the genome duplication to have occurred after the divergence of AG1-IA from AG1-IB (Fig. 2). This hypothesis has been corroborated by the observation that, out of 669 paralogous gene pairs in AG1-IA, 620 gene pairs have orthologs in the AG1-IB genome assembly and 547 (88%) of those are present in single copy (only one ortholog) in AG1-IB genome (Table S9).

### Transposable elements are associated with genome duplication and evolution

Transposable elements (TEs) are abundant in filamentous fungi^25^. They have been anticipated to create localized blocks in the duplicated genomic regions by inserting breakpoints and increasing rates of chromosomal rearrangements thereby creating genomic variation^25, 26^. We noticed the presence of TEs within sixty-two paralogous blocks, amongst the total 92 paralogous blocks (Fig. 3a). The genome-wide comparison of the density of the TE revealed an overall average of 1120 bp of TE per 10 kb of genome while, the density of TE within the duplicated blocks was 1461 bp per 10 kb and the TE density within the 10 kb flanking regions of the duplicated blocks was 1665 bp per 10 kb. The appearance of high frequency of TEs in the duplicated block has introduced disruption in the continuity of the synteny. Merging the duplicated blocks that are within 200 kb after discounting the TEs would have extended the duplicated blocks from 15% to 22.1% of the genome assembly. Together, these analyses suggest that whole genome duplication and TE-mediated gene disruption have a profound effect on *R. solani* AG1-IA genome evolution.

**Fig. 3.**
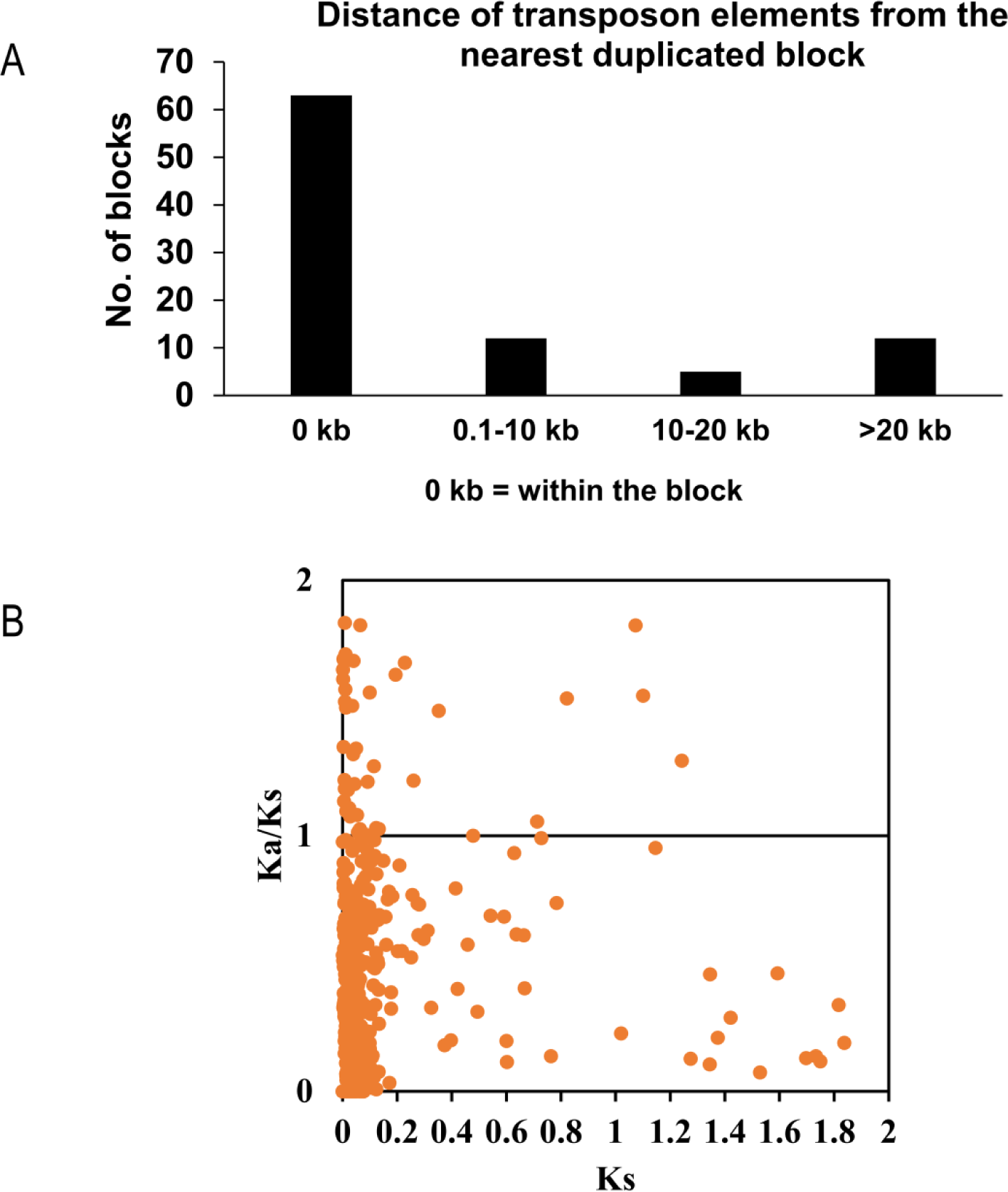
Distribution of TEs in *R. solani* genome and neofunctionalization of *R. solani* genes. A. Distance of the transposon elements nearest to a duplicated paralogous gene block. Number of paralogous gene blocks were plotted against the distance of the nearest transposon elements. 0 kb denotes where the transposon elements reside within the paralogous blocks. B. A scatter plot of the ratio of the rates of non-synonymous to synonymous substitutions against the synonymous substitution of the paralogous gene pairs of *R. solani* AG1-IA.

### Neo-functionalization of genes due to gene duplication event

To determine whether the gene duplication has led to the emergence of new functions, we calculated ratio of non-synonymous substitution rate to synonymous substitution rate (Ka/Ks) of the TE-associated paralogs and plotted against the synonymous substitution rate (Ks). Sixty-four gene pairs were identified to have Ka/Ks >1, suggesting neo-functionalization after gene duplication (Fig. 3b, Table S10). We identified six paralogous pairs wherein one paralog encodes a potentially secreted protein whereas, the other paralog encodes a non-secreted protein (Table S11). In one of such paralog pairs (Rs_09385-Rs_11744), the paralog (Rs_09385) with secretion signal showed significantly higher expression during pathogenesis in rice, compared to the paralog (Rs_11744) that lacks the secretion signal (Fig. S7). This suggests *R. solani* to have adapted one of the duplicated paralog as secreted effector to facilitate host colonization. However, in two other cases (Rs_07297-Rs_11800 and Rs_11307- Rs_11399), the paralog with secretion signal exhibits relatively less expression during pathogenesis in rice, compared to the paralog without secretion signal (Fig. S7). It seems *R. solani* downregulates the expression of the secreted paralogs to potentially avoid host recognition and induction of defense response.

Out of 669 duplicated gene pairs, we observed 50 pairs to possess at least one different domain altogether and 107 pairs to have a disrupted/missing domain in one of the partners (Table S12). Notably, 31 of the duplicated paralogous gene pairs exhibited loss of functional domain in one of the paralogs. For one of the pairs (Rs_06191-Rs_10629), we observed the paralog with additional domain (Rs_06191; Glycosyl hydrolase) had significantly higher expression during establishment phase (1 dpi) of pathogenesis in rice, compared to the domain deleted paralog (Rs_10629) (Fig. 4a). However in case of Rs_09094-Rs_11334 duplicated pair, the paralog with domain deletion (Rs_11334) exhibited significantly higher expression during necrotrophic (2-3 dpi) phase of *R. solani* pathogenesis in rice, compared to the one (Rs_09094; GMC oxidoreductase) with additional domain (Fig. 4b). Differences in the expression of the duplicated gene pairs of *R. solani* during colonization in rice highlighted neo-functionalization. In order to analyse the importance of the genes undergoing neofunctionalization in *R. solani*, we used a dsRNA based approach to silence *R. solani* genes and study their importance during pathogenesis in tomato^21^. We used gene-specific dsRNA to downregulate two duplicated paralogous gene pairs (*Rs_06191*-*Rs_10629* and *Rs_09094*-*Rs_11334*) of *R. solani* to study the impact of gene silencing during pathogenesis. The qRT-PCR analysis reflected efficient silencing of each of the target genes in dsRNA treated *R. solani* mycelia (Fig. S8a). Notably, silencing of *Rs_06191* but not *Rs_10629*, whereas silencing of *Rs_11334* but not *Rs_09094* compromised the pathogenesis of *R. solani* (Fig. 4c-e). The disease symptoms (Fig. 4c), disease severity index (Fig. 4d) and pathogen load (Fig. 4e) were significantly reduced in plants infected with *Rs_06191* and *Rs_11334* silenced *R. solani* mycelia, compared to the control plants, infected with buffer-treated *R. solani* mycelia. On the other hand, silencing of *Rs_10629, Rs_09094* and a previously reported negative control gene *Rs_GT34* (Glycosyl transferase family protein 34)^21^, did not compromise the pathogenesis of *R. solani* (Fig. 4c-e).

**Fig. 4.**
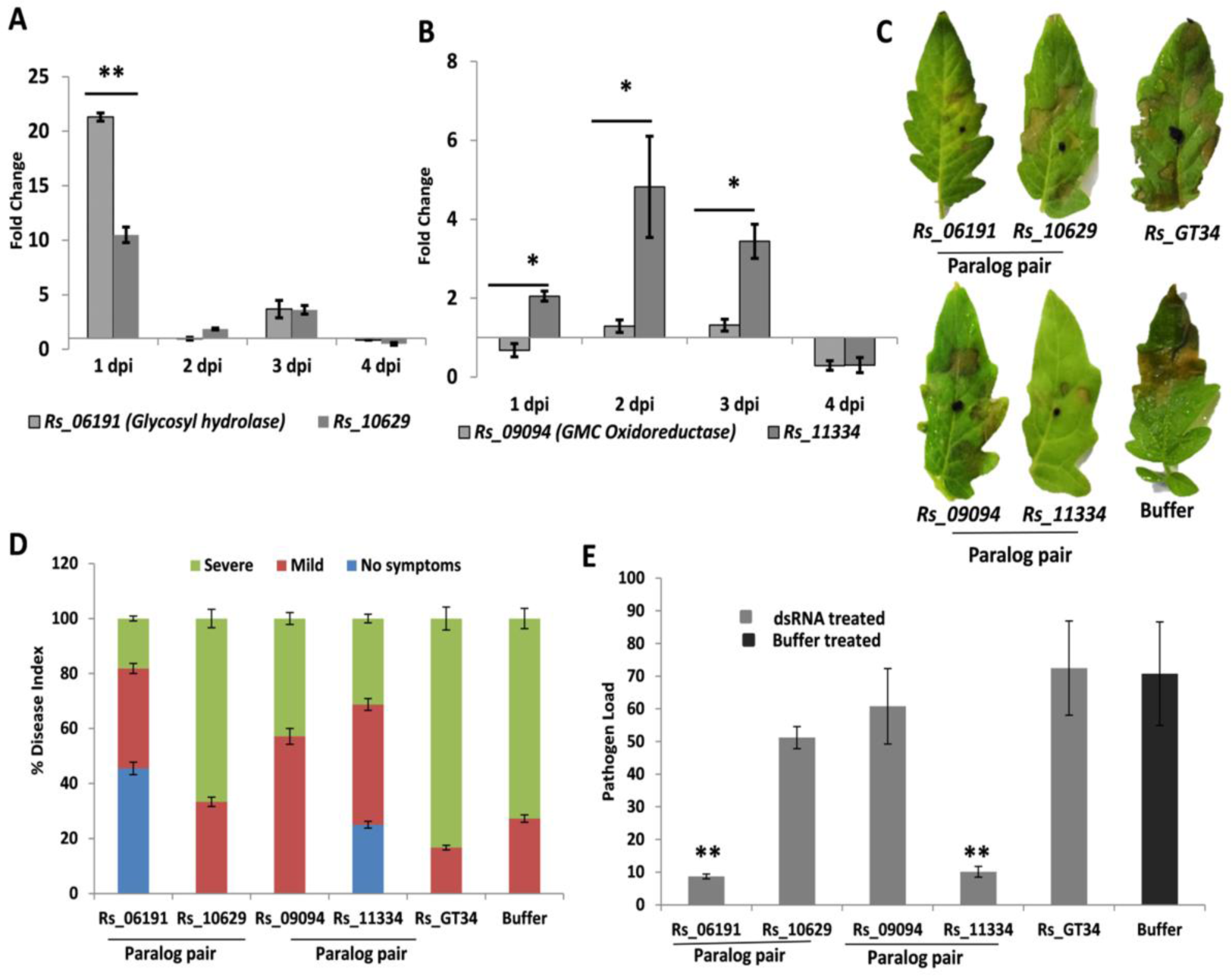
Loss and gain of domain in paralog gene pairs of *R. solani* AG1-IA modulate its pathogenesis in tomato. qRT-PCR based expression analysis; 2^−ΔΔCt^ of A. paralog pair (Rs_06191; with glycosyl hydrolase domain and Rs_10629; lacking the domain) and B. Paralog pair (Rs_09094; with GMC oxidoreductase domain and Rs_11334; lacking the domain) upon *R. solani* infection in rice, at different time points. The fold change in gene expression was estimated with respect to 0 dpi, using *18S rRNA* of *R. solani* for normalization. C. Disease symptoms, D. Disease index and E. Pathogen load (2^−ΔCt^ abundance of 18S rRNA of *R. solani*, using *actin* gene of tomato for normalization) in leaves infected with gene silenced (dsRNA treated) and buffer treated (control) *R. solani* mycelia, at 3 dpi. The *Rs_GT34* was used as negative control. Graph shows mean values ± standard error of three biological replicates. (**) indicate significant difference at p ≤ 0.01 and (*) indicate significant difference at p ≤ 0.05. glucose–methanol–choline oxidoreductases.

### Molecular diversity and population genetic structure of the Indian isolates of *R. solani* **AG1-IA**

To understand the genetic diversity, forty-two isolates of *R. solani* AG1-IA collected from rice fields of six geographical regions (four directional zones, north-east and the central) of India (Table S13) were sequenced with 350x throughput to produce 700.853 Gb sequence data. 5,046,121 SNPs were obtained by physically mapping the reads on the presently assembled 74 genomic contigs of *R. solani* AG1-IA strain BRS1 and a wide range of distance coefficient from 0.00008 to 0.14 with an average of 0.06 was observed. A principle component analysis (PCA) based on the genomic distance distributed the Indian isolates in three different groups and a subgroup of admixture between group I and group II, suggesting natural hybridization amongst them. More than 50% (n=22) of these isolates belonged to group II (Fig. 5a).

**Fig. 5.**
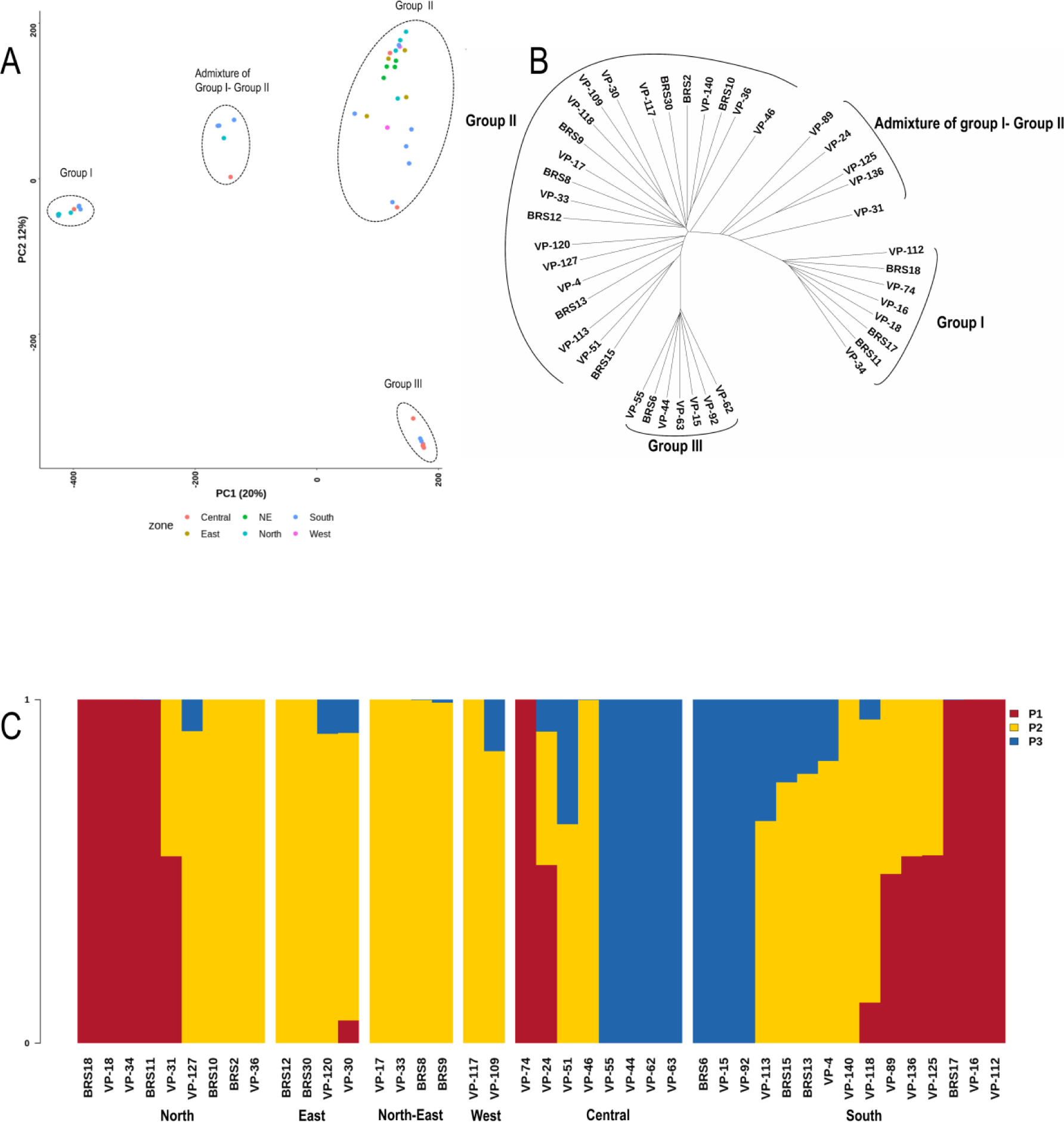
Classification of the Indian field isolates of *R. solani* AG1-IA based on their genomic diversity. A. Principle component analysis of 42 Indian AG1IA isolates clustered them into 3 major groups and an admixture group. B. Unrooted dendrogram depicting the genetic relationship among the isolates. C. Population structure inferred by ADMIXTURE analysis at K=3.

There was no correlation between the genetic distance and the geographic locations from where those were isolated. The groups are genetically equidistant from each other. The minimum distance coefficient between the group I and group II is 0.172, and that between the group II and group III, and between the group I and group III are 0.168 and 0.17, respectively. The genetic relationship among the forty-two isolates is depicted in an unrooted dendrogram (Fig. 5b). The SNPs clearly discriminated all the 42 isolates from each other resulting in definite clusters as observed in the PCA. The population genetic structure was determined among these isolates using the SNPs by optimizing the K value at 3 and the genomic compositions of the isolates were presented in three K colour segments. Twenty-six isolates in these three groups showed pure genetic background (Fig. 5c). All the four isolates collected from the North-East regions of India showed pure genomic background probably due to geographic distance from the other parts of India. Most of the other isolates showed hybridization of two groups. All the group III isolates (n=7) were collected from two neighboring states Chhattisgarh and Telangana and they showed pure genetic background.

In order to understand genetic diversity within the Indian isolates of *R. solani* AG1-IA, we calculated the average pairwise nucleotide diversity within population (θπ), Waterson’s estimator of segregating sites (θ⍵) and Tajima’s D, the commonly used metrics of genetic diversity^2728^. Plotting diversity metrics in sliding windows across the genome revealed a high θπ all over the genome however, high θ⍵ at some of the TE-rich regions (Fig. 6). A total 1028 genomic regions with Tajima’s D values more than 2 (representing 1588 genes) were predicted to be the targets of diversifying selection. They encode 288 PHI-base homologs, 34 CAZymes, 19 putative effectors and 83 putative secreted proteins. On the other hand, sixteen regions (representing 18 genes) with Tajima’s D values less than 2 were predicted to be the targets of purifying selections and they are located in the TE-rich regions (Fig. 6, Table S14).

**Fig. 6.**
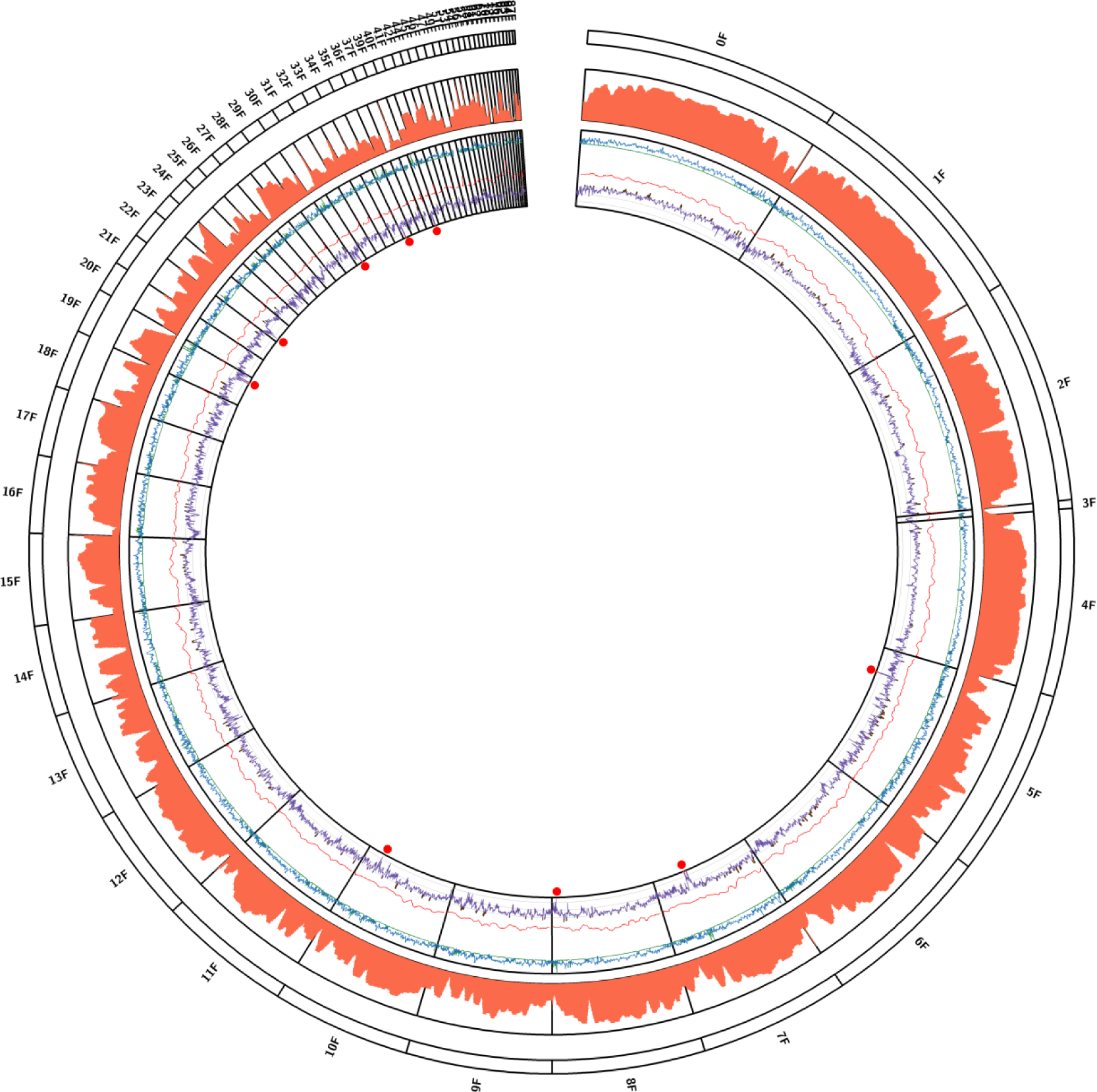
Genomic variations in the Indian isolates of *R. solani* AG1-IA and identification of genomic regions of high and low variations. Diversity metrics, presented as average pair- wise nucleotide diversity θπ (Blue line, min=0, max=0.5), θ⍵ (Green line, min=0, max=0,5), FST (Red line, min=0, max=1.0) and Tajima’s D (Purple line, min=-3, max=+3). The regions with increased (>+2) Tajima’s D denoting positive selection and with decreased (<-2) Tajima’s D denoting purifying selection. Regions denoting purified selection are marked by red dots. The gene density plotted in orange colour. Gene density and FST are plotted In 100 kb sliding window. The other values are plotted in 500-SNP sliding window with step size 100-SNP using Circos.

Further, we studied the expression dynamics of some of the genes under positive and purifying selection, during pathogenesis in rice. We observed 14 genes (including three LPMO_AA9 (Lytic polysaccharide monooxygenase_Auxillary activity family 9 domain containing proteins) under positive selection to be upregulated during 2-3 dpi of *R. solani* infection in rice (Fig. 7a, Supplementary Fig. S9), which coincides with the transition from establishment (biotrophy) to necrotrophy phase^29^. Notably, dsRNA mediated silencing of the gene encoding one of the GH61/LPMO_AA9 domain containing protein (Rs_01468) compromised the pathogenesis of *R. solani* in tomato (Fig. 7). The disease symptoms (Fig. 7b), disease severity index (Fig. 7c) and pathogen load (Fig. 7d) were significantly reduced in plants infected with *Rs_01468* silenced *R. solani* mycelia, compared to those infected with buffer treated (control) or *Rs_GT34* silenced mycelia. Moreover, qRT-PCR analysis reflected dsRNA mediated efficient silencing of the target gene during pathogenesis (3 dpi) in tomato (Fig. S8b).

**Fig. 7.**
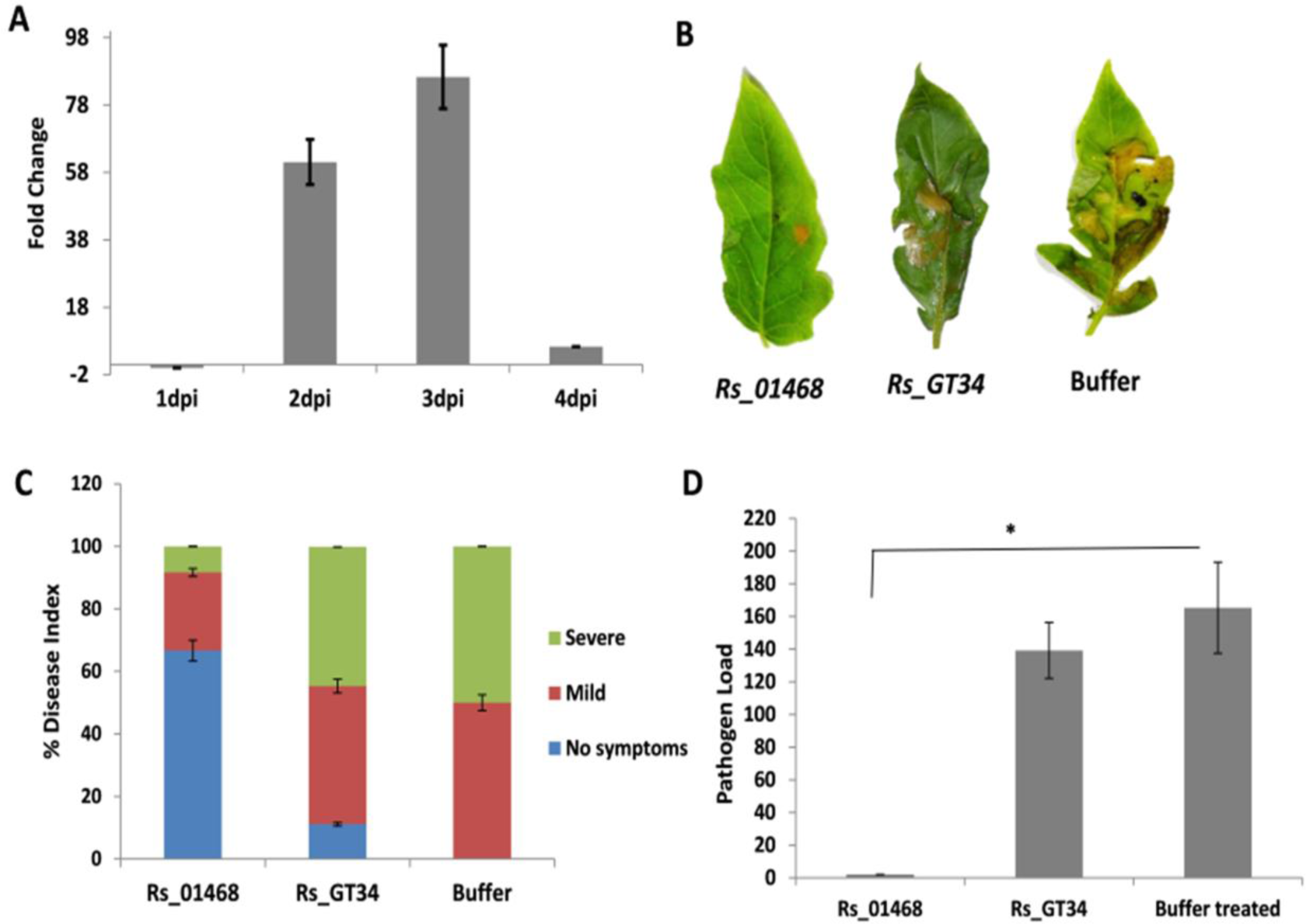
Silencing of positive selection gene *Rs_01468* (LPMO) compromises pathogenesis of *R. solani*. A. qRT-PCR based expression analysis of *Rs_01468* during pathogenesis of *R. solani* in rice at different time points compared to 0 dpi. B. Disease symptoms, C. Disease index and D. Pathogen load in tomato infected with *Rs_01468* silenced *R. solani* compared to control (*Rs_GT34* silenced and buffer treated), at 3 dpi. *R. solani* 18S rRNA or tomato actin gene was used as endogenous control. Graph shows mean values ± standard error of three biological replicates. (*) indicate significant difference at p ≤ 0.01. LPMO = Lytic polysaccharide monooxygenase, GT34 = Glycosyl transferase family 34 protein.

Amongst genes undergoing purifying selection, the *Rs_11537* encoding glucosamine phosphate N-acetyltransferase (GNAT) was upregulated during 2-3 dpi of *R. solani* pathogenesis in rice (Fig. 8a). The *R. solani* strains encode large number of GNAT proteins with size ranging from ∼80-700 aa. Notably Rs_11537 is only 80 aa long and phylogenetic analysis reflected it to be in a clade with a few other smaller size GNAT proteins (two of them were 87 aa long while Rs_05222 and some others were ∼183 aa) (Fig. S10). This tempted us to speculate that the Indian strain of *R. solani* (BRS1) has particularly selected the smaller sized GNAT protein to effectively colonize plants. The qRT-PCR based gene analysis reflected that expression of smaller sized (*Rs_11537*) but not the bigger sized (*Rs_05222*) GNAT was induced during pathogenesis of *R. solani* in rice (Fig. 8a). Further using the dsRNA approach we downregulated the *Rs_11537* and *Rs_05222* genes and studied the impact on pathogenesis of *R. solani* in tomato. The qRT-PCR analysis reflected efficient silencing of the target genes (Fig. 8b). Moreover, no cross-silencing of *Rs_05222* gene was observed upon dsRNA mediated silencing of *Rs_11537* (Fig. 8c). Notably, plants infected with *Rs_11537* silenced *R. solani* mycelia had significantly reduced necrotic disease lesion (Fig. 8d), disease index (Fig. 8e) as well as pathogen load (Fig. 8f), compared to those infected with *Rs_05222*/*Rs_GT34* silenced or buffer treated mycelia.

**Fig. 8.**
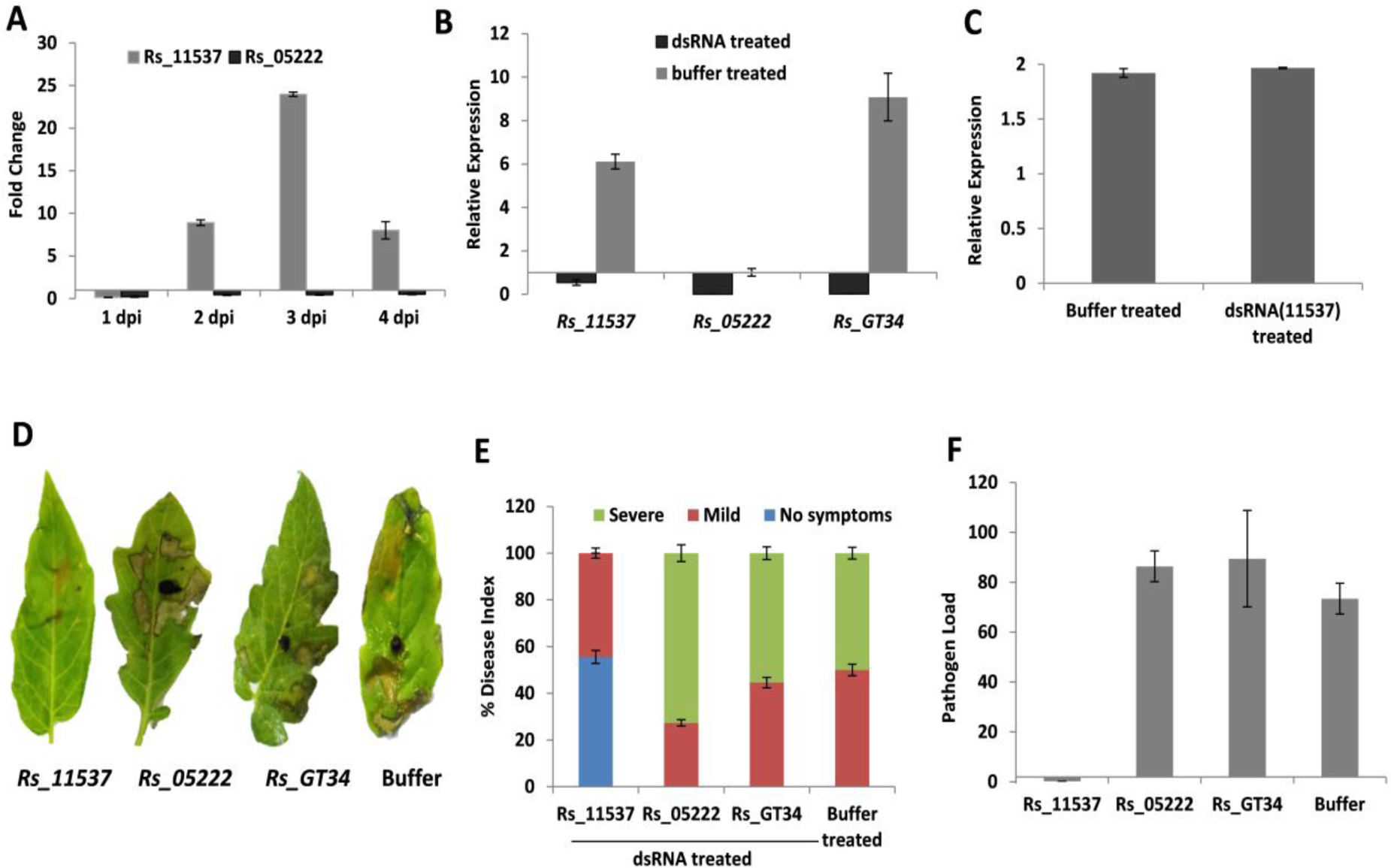
*Rs_11537* (GNAT), a gene under purifying selection is important for pathogenesis of *R. solani* in tomato. **A)** qRT-PCR based expression analysis of *Rs_11537* and *Rs_05222* genes during *R. solani* pathogenesis in rice at different time points as compared to 0 dpi using rice actin gene for normalization. qRT-PCR based expression analysis of *R. solani* genes **B)** *Rs_11537*, *Rs_05222* and *Rs_GT34* genes during 3dpi of pathogenesis in tomato infected with dsRNA treated and buffer treated mycelia. **C)** qRT-PCR based expression of *Rs_05222* in tomato infected with *Rs_11537* silenced *R. solani.*. **D)** Disease symptoms, **E)** Disease index and **F)** Pathogen load in tomato infected with *Rs 11537*, *Rs_05222* and *Rs_GT34* silenced *R. solani* compared to buffer treated, at 3 dpi. Graph shows mean values ± standard error of three biological replicates.

## Discussion

The *Rhizoctonia solani* is an important fungal model system to study the genetic adaptation of the pathogens to colonize a wide range of plant species. In this context, it is important to not only obtain a better genome assembly but also analyze the genetic diversity amongst pathogenic isolates of a particular anastomosis groups (AGs) or between different AGs of *R. solani*. In this study, utilizing hybrid genome assembly of Pacbio and Illumina reads, we have not only produced so far the best genome assembly of an AG1-IA strain genome and identified a large number of unigenes, but also have shed important insights about diversification of different AGs. The synonymous nucleotide substitution rates within the *R. solani* anastomosis groups suggests that diversification of *R. solani* strains into various AGs have happended over the last ∼44 million years. Similarly, most of the extant crop plants have undergone whole genome duplication and diversification over the last 60-66 million years (during the tertiary period of Cenozoic era) to adapt to different ecological niches^30^. Our data suggests that around similar period, *R. solani* strains have diversified into different AGs not only to colonize different plant species (members of Solanaceae, Asteraceae, Poaceae) but have also specialized to infect different plant parts (such as roots, stem and leaves). Notably, the AG3 and AG2- 2IIIB strains, which diverged from the other AGs ∼44 mya predominantly infect underground plant parts whereas AG1-IA, AG1-IB and AG8, which are closely related and diverged over the last ∼32 million years infect aerial plant parts such as stem and leaves^11, 31, 32^.

Evidence of whole genome duplication followed by gene family expansion by gene gain and loss has been reported in fungal pathogens^26, 33, 34^. This has been proposed to cause diversification of host range and increasing virulence. Previous genome based studies had revealed expansion/emergance of gene families/orthogoups in different AGs of *R. solani*^8, 14^. We have observed similar expansion of gene families in AG1-IA in comparison to its closest relative AG1-IB especially, the genes located in the duplicated gene blocks. About fifteen percent of the *R. solani* AG1-IA strain BRS1 genome encompasses the duplicated region. Moreover, we identified large number of paralogous gene pairs, grouped into 46 pairs of duplicated syntenic blocks (each having 5 to 58 duplicated genes). We emphasize that whole genome duplication followed by possible sequential duplication events have contributed towards evolution of AG1-IA strains. Our data suggests that the gene duplication has occurred in AG1-IA after divergence from AG1-IB. Generally, the processes of gene duplication/gene family expansion have been anticipated to enable the organism to adapt to changing environmental conditions. For example, the gene expansion events have enabled *Rhizopus oryzae* (causal organism of mucormycosis) to utilize complex carbohydrates and rapidly generate energy to ensure rapid growth^26^. We observed genes encoding CAZymes, PHIbase, secreted/effectors, transcription factors, and secondary metabolite-related proteins to have duplicated in *R. solani*. The expansion of pathogenicity associated genes/families suppots that *R. solani* is equipped for adopting diverse strategies to colonize broad range of plant species.

It is to be noted that natutal infection by AG1-IA strains of *R. solani* are more prominently reported than other AGs^35, 36^. Presence of extensive amount of interspersed repeat elements has introduced several brakes in the duplicated gene blocks, due to which the probability of frequent recombination and gene loss as well as modification due to domain loss has increased in *R. solani*. We anticipate that TE mediated gene duplication events and subsequent gain/loss of functional domain has enabled the *R. solani* to adopt newer functions. Indeed, we observed altered gene expression patterns of the paralogous gene pairs during infection process, which suggests neofunctionalization of the duplicated genes^37^. Moreover, our data that silencing of one of the paralogous pair but not the other pair compromises the pathogenesis of *R. solani*, further emphasize that TE-mediated disruption of genes or domain loss has impact on pathogenesis of AG1-IA.

Generally, there is constant arms race between the plants and the pathogens, to prevent and establish disease, respectively. The plants evolve mechanisms to detect the pathogens and mount potent defense response to ward-off them. On the otherhand, pathogens rapidly evolve strategies to avoid host recognition and/or suppress host defense response. Considering the genes undergoing positive selection and loss of functional domains/secretion signal in one of the genes of the paralogous pairs, we anticipate *R. solani* to have adopted new functions for several of them. Notably, the expression of certain paralogs with secretion signal or additional domain were significantly less during pathogenesis in rice, compared to the corresponding paralogs lacking secretion signal or additional domain. This emphasizes that *R. solani* may have evolved to downregulate the expression of such paralogs to potentially avoid host recognition. On the otherhand, in certain other cases, the paralogs with secretion signal or additional domain showed higher expression during pathogenesis, compared to others that lack secretion signal or additional domain. This emphasises such paralogs to be involved in promoting successful host colonization.

To avoid host recognitions pathogens try to evolve newer alleles of genes that are important for pathogenesis. The estimation of average pairwise nucleotide diversity index (Tajima’s D) amongst different pathogenic field isolate of *R. solani* has enabled us to identify genes under positive selection. Positive or diversifying selection refers to the Darwinian theory of natural selection, wherein new alleles/genes emerge as well as change rapidly for adaptation to the external environment. In our study, we observed several genes encoding effectors, cell wall degrading enzymes and pathogenicity determinants (PHI-base homologs) to be under positive selection. The induced expression of several of them during pathogenesis of *R. solani* in rice emphasize their importance during pathogenicity process. Notably, silencing of one of them i.e *Rs_01468* (encoding GH61, LPMO_AA9 domain containing protein) significantly compromised pathogenesis of *R. solani*. The LPMO domain containing proteins are widespread in filamentous fungi including *R. solani* and enables the pathogens to degrade different components of the host cell wall^8, 38^. In *Colletotrichum*, induction of LPMO has been associated with switching between biotrophic and necrotrophic phases^39^. Considering that *Rs_01468* is induced during 2 and 3 dpi of *R. solani* pathogenesis, we anticipate that it may facilitate transition of biotrophy to necrotrophic phase in *R. solani*.

We observed 18 genes to be under purifying selection in *R. solani* and they encode important functions such as reverse transcriptase (Rs_06070; Rs_06071), DNA polymerase (Rs_11453), Sir2 family transcriptional regulator (Rs_11455) and glucosamine-phosphate N-acetyltransferase (Rs_11537). Considering the importance of Glucosamine 6-phosphate N-acetyltransferase (GNAT) domain containing proteins in biosynthesis of chitin in the fungal cell wall ^40, 41^, we investigated the involvement of *Rs_11537* in pathogenesis of *R. solani*. The gene was upregulated during 2-3 dpi of *R. solani* pathogenesis that coincides with the transition of biotrophy to necrotrophic phase. The dsRNA mediated silencing of the gene effectively compromised the pathogenesis of *R. solani*. It is to be noted that homologs of GNAT genes are abundantly present in *R. solani* and relatively shorter sized homologs (80-87 aa) are under purifying selection. Indeed, silencing of a larger sized GNAT homolog (Rs_05222; 183aa) does not compromise the pathogenesis of *R. solani*.

Genome analysis of Indian isolates of *R. solani* AG1-IA revealed presence of three distinct clades and one admixture of group I and II. We did not observe any correlation between the geographic distribution or virulence of these isolates and their genome sequences. Existence of five isolates as admixtures indicates natural exchange of genetic materials between the *R. solani* isolates. It is to be noted that horizontal transfer of lineage-specific genomic regions between the fungal strains at a level of even one-quarter of the whole genome and acquisition of new property has been reported before^42^.

In summary, a high quality genome assembly of *R. solani* AG1-IA enabled us to find out a recent whole genome duplication followed by transposon element-mediated gene loss that shaped the present genomic structure of this agriculturally important rice pathogen. This has caused an expansion and domain alterations of the gene families associated with it’s virulence. Genome-wide analysis of multiple isolates identified genomic regions which are essential for the survival and therefore, are not allowing any change. The analysis has also identified the regions that are continuously evolving and being positively selected by nature enabling the pathogen to adopt to the changing environment and maintain pathogenicity.

## Materials and Methods

### Biological materials

Different strains of *Rhizoctonia solani* AG1-IA were grown on Potato Dextrose Agar (PDA; 39 g/L; Himedia, Mumbai, India) plates at 28°C. The growth rate, maturation of sclerotia, sclerotial size and number etc of *R. solani* strains were measured, as described earlier^21^. Also the pathological attributes of these strains were studied in rice (indica cultivar PB1) and relative vertical sheath colonization (RVSC) was calculated at 3 dpi (days post inoculation) as described earlier^14^.

### Genome sequencing and assembly

High-molecular weight DNA was extracted from *R*. *solani* AG1-IA strain BRS1, as described earlier^7^ and genomic DNA fragment library was constructed for PacBio SMRT (single molecule real time) sequencing, as per the manufacturer’s instructions. A total of 13.74 Gb (∼300x) sequence data was generated from two PacBio Sequel runs. Further, two Illumina libraries were prepared with 2 kb and 5 kb insert sizes and a total of 2.99 Gb (2 × 150 base pairs) (∼67x) sequence data with 2 kb insert size and 3.1-Gb (2 × 150 base pairs) (∼69x) sequence data with 5 kb insert size were generated.

The *de novo* assembly of the genome was performed using FALCON and FALCON-Unzip (pb-falcon 0.2.7)^43^ tools. FALCON-Unzip tool assembles a set of partially phased primary contigs and fully phased haplotigs. The initial assembly with FALCON was carried out with parameters set, 50 Mb for genome size, 30x for seed coverage and 1000 bp as the length cut- off for seed reads. Further, FALCON-Unzip module was applied to phase the raw reads according to the SNPs identified in the FALCON assembly and reassemble in discrete haplotype specific manner. The genome assembly was polished and consensus sequences were attained with Arrow polishing tool in FALCON-Unzip. The Illumina pair-end reads with 5 kb and 2kb insert size were used for sequence correction using Pilon v1.23^44^. The reads were mapped to the polished assembly using BWA v0.7.17^45^. Samtools v1.9^46^ along with Pilon were run with parameters “–diploid –fix all” to correct bases, fix mis-assemblies and fill gaps.

### Genome annotation

The repetitive sequences, identified by RepeatModeller v2.0.1^47^ and Repbase19 database^48^, were used to mask *R. solani* genome with RepeatMasker v4.1.0 (http://www.repeatmasker.org/). Three different approaches i.e. *ab initio*, homology-based and transcript-based prediction were used to predict the protein-coding genes in the Repeat-masked genome assembly. AUGUSTUS v3.3.3^49^ with parameters trained on Coprinus species and GeneMark-ES v4.59^50^ with training data customized for fungus were used for *ab initio* prediction. For homology-based gene prediction, we used EXONERATE v2.2.0^51^ and AAT r03052011^52^ tools and protein sequences predicted from AG1-IA, AG8 draft assemblies. In transcript-based approach, we aligned AG1-IA transcriptome sequences for spliced alignment using PASA v2.4.1^53^. The predictions from these three approaches was integrated using EVIDENCEMODELLER (EVM) v1.1.1^54^ to generate consensus gene models. Finally, for the identification of spliced variants and prediction of untranslated regions the EVM output was run through PASA.

Calculation for the probability of sequential genome duplication was done following the method reported earlier^17^. In case the duplicated regions are created in a sequential manner, those will follow a Poisson distribution in the genome with the formula *f(x;λ)* = *λ^x^. e^-λ^/x*!, where e = 2.71828, x is the probability of which is given by the function and is a positive real number equal to the expected number of occurrences that occur during the iven interval. According to this equation, we expect 18.4 triplications per 100 duplicates. Instead of expected 8 triplication out of 46 duplicates, we have observed 4 triplications. The probability for this observation is *p*(4;8) = 0.057.

### Gene family classification

In order to predict the gene family, proteins derived from genome assemblies of multiple *R. solani* strains belonging to AG1-IA (GCA_000334115.1, GCA_015342405.1, GCA_015342435.1, GCA_015341985.1, GCA_015342415.1, BRS1), AG1IB (GCA_000832345.2, GCA_000350255.1), AG3 (GCA_000715385.1, GCA_000524645.1), AG8 (GCA_000695385.1) and AG2-2IIIB (GCA_001286725.1) were included in the analysis. The full set of proteins for each strain was used to infer gene family (orthogroups) with OrthoFinder v2.4.0^55^. DIAMOND blast with E-value<1e-05 and MCL clustering algorithm was used to identify similarity index and cluster them.

### Functional annotation

Sub-cellular localization, secretion status and transmembrane domains were predicted using Phobius v1.01^56^, SignalP v5.0^57^, TMHMM^58^ and TargetP^59^ tools. The bigPI Fungal Predictor ^60^ was used to identify GPI modification sites. The secondary metabolite encoding genes were identified using AntiSMASH fungal version^61^. The putative genes involved in pathogen-host interactions were predicted based on sequence similarity (*E-*value <10^−^5) in PHIbase (Pathogen Host Interactions Database)^62^. InterProScan v5.28-67.0^63^ was used to assign GO terms and identify conserved domains including fungal specific transcription factors. The CAZymes encoding genes were predicted using dbCAN2 meta server (http://bcb.unl.edu/dbCAN2/) by HMMER, DIAMOND and Hotpep based searches. Effectors were predicted using SignalP v5.0 and EffectorP v2.

### Evolutionary analysis

Syntenic blocks within AG1-IA strains and between AG1-IA and other anastomosis group members were identified using MCScanX^64^ with default parameters and synteny distribution was plotted with Circos^65^. Synonymous (*Ks*) and nonsynonymous (*Ka*) substitutions rates of homologous gene pairs were calculated with the perl script, add_ka_and_ks_to_colinearity.pl incorporated in MCScanX. Median *Ks* value for each block was considered to be representative of the duplicated region. *Ks* distribution were plotted to estimate divergence times and genome duplication events. The time was calculated at the peak Ks value using the formula T=Ks/2*r,* where *r* is the fungal neutral substitution rate (r=1.3x10^-8^)^66^. Expected number of triplications and the probability of observed triplications is calculated as per the methods described previously^17^.

### SNP genotyping and population structure of Indian AG1-IA strains

The high-molecular weight DNA of different *R*. *solani* strains isolated from different parts of India (Table S13) was extracted, as described earlier^7^ and subjected to Illumina (Novaseq) sequencing (2 X 150 bp), as per the manufacturer’s instructions. A total of 4 Gb (∼10x) sequence data was generated for each of the strains and low quality reads (PHRED score < 20 and length < 30 bp) were trimmed. The trimmed reads were mapped to AG1IA reference genome with BWA v0.7.17-r1188 and analysed for PCR duplicates with MarkDuplicates in Picard v2.23.3 (https://broadinstitute.github.io/picard/). The SAM files were sorted, indexed and converted to BAM format with Samtools v1.9. Using Genome Analysis Toolkit (GATK v4.1.8.1^67^), variants were initially identified by HaplotypeCaller with option -ERC GVCF and then combined genotyping was performed with GenotypeGVCFs. SNP variants were selected with SelectVariants and filtered for QD < 2.0, FS > 60.0, MQ < 40.0, MQRankSum < -12.5, ReadPosRankSum < -8.0 and SOR > 4.0. The variants were annotated by SnpEff ^68^ based on the annotation GFF file. PCA (principal component analysis) was conducted with TASSEL v5^69^ and the result was plotted with ggplot. A phylogenetic neighbor-joining tree was generated from the numerical genotype data using TASSEL and visualized with iTOL (interactive tree of life https://itol.embl.de/). The commonly used diversity matrices such as Pi, Theta, Fst and Tajima’s D were also calculated with TASSEL with default parameters. The population structure was determined by ADMIXTURE v1.3.0^70^ with K varying from 2 to 5 and K=3 had the lowest cross validation error and the ancestry distribution of each sample was illustrated with R script.

### Development of *R*. *solani* genome database

A Rice sheath blight (RSB) database has been developed to host the genome assembly and annotation of *R*. *solani* AG1-IA genome. The database operates on a linux system and can be assessed by the URL: https://nipgr.ac.in/RSB/. The current database framework is built on the Apache server and the RSB web interface has been designed with laravel (https://laravel.com/), an open-source PHP framework, HTML (https://html.spec.whatwg.org/), CSS (https://www.w3.org/Style/CSS/Overview.en.html), and JavaScript (https://www.javascript.com/). In addition, RSB is integrated with stand-alone BLAST^71^ for online similarity search against genome, proteome, CDS or mRNA of any anastomosis groups of *R*. *solani*, jBrowse^72^ for interactive visualization of the genome, and batch download for genome, mRNA, CDS, and protein.

### Phylogenetic analysis

Annotated GNAT proteins in *R. solani* were identified and extracted from NCBI and used to find members in AG1IA assembly based on homology. The sequences were examined for the GNAT family domains and classified accordingly. Multiple sequence alignment of the protein sequences was performed using CLUSTALW. The neighbour joining tree with 1000 bootstraps was constructed with MEGA X software using p-distance model and pairwise gap deletion.

### Pathological assays

For infection in rice (indica cultivar PB1), freshly grown equal sized *R. solani* strain BRS1 sclerotia were inoculated within the sheaths of ∼45 old rice plants. The disease symptoms were quantified in terms of relative vertical sheath colonization as described earlier ^14^. The infected tissues (including 1 cm up and down from the site of infection) were further harvested for expression analysis at 0, 1, 2, 3 and 4 dpi. A total of 3-4 sheaths per plant were infected and total 4-5 plants were used for the experiment.

In tomato; the *R. solani* sclerotia were attached to abaxial surface of the leaves of 30 day old tomato plants (*Solanum lycopersicum*; Pusa Ruby) using aluminium strips^73^. The plants were further incubated in growth chamber at 26°C temperature under 80% relative humidity and 12/12hr of day/night cycle. The disease symptoms were recorded at 3 dpi and disease severity was categorized into severe, medium, negligible symptoms. A total of 4-5 plants and three leaves per plants were infected with *R. solani* and each experiment was repeated three times.

### qRT-PCR based expression analysis and pathogen load quantification

The qRT-PCR-based expression analysis of *R. solani* genes was carried out during pathogenesis in rice (cv. PB1) and tomato (cv. Pusa Ruby). The primers were designed to selectively amplify the target pathogen genes (Table S15) and qRT-PCR was performed using Sso Advanced Universal SYBR Green Supermix (BioRad) according to manufacturer’s instructions. The relative expression was calculated using 2^-ΔCt^ method, wherein ΔCt is the difference between Ct values of target and reference (*18S rRNA*) genes of *R. solani*^74^. The fold change was calculated using 2^-ΔΔCt^ method^75^ wherein ^ΔΔ^Ct is the difference in the expression of *R. solani* genes at 1 dpi, 2 dpi, 3 dpi and 4 dpi, compared to 0 dpi. The fungal load was quantified in the infected samples, using 2^-ΔCt^ method, wherein ΔCt is the difference between Ct values of fungal *18S rRNA* (target) and host *actin* gene (as reference), as described earlier^73^. The data from three independent biological replicates were used to calculate standard error and one-way ANOVA was performed using Sigma Plot 11.0 (SPSS, Inc. Chicago, IL, USA) software to estimate the statistical significance (determined at *p≤ 0.05*) between separate groups using Student-Newman-Keuls Test.

### dsRNA mediated silencing of *R. solani* genes and functional studies

For dsRNA mediated silencing of *R. solani* genes, the sequences were analysed using siFi21 software (http://labtools.ipk-gatersleben.de/)76 to select unique fragment that have no off-target silencing effect in *R. solani* as well as host (tomato) genome. The target regions were PCR amplified from the *R. solani* cDNA using gene specific primer pairs having T7 promoter sequence at 5’ end of the forward primer (Table S15). For in-vitro transcription, 1µg of the purified gene fragments was used to produce dsRNA, as per the manufacturer’s protocol (using MEGAscript T7 Transcription Kit; Thermo Scientific). The amount of dsRNA synthesized was measured using Nanodrop (Themo Scientific) and 50 µg of dsRNA was used to treat *R. solani* sclerotia. The sclerotia were grown on 1% agar plates and the mycelial mats were treated with dsRNA targeting particular genes for 24 hrs at 28°C, and the silencing of the gene was investigated using qRT-PCR, following the protocol described earlier^21^. The list of primers used in the study is enlisted in Table S15.

To study the effect of gene silencing on fungal growth in laboratory media, the dsRNA treated mycelia were placed on 1% agar plates and incubated at 28^0^C. The mycelial growth was quantified in terms of diameter of the mycelial spread on the agar plate. Further to investigate the effect of gene silencing on pathogenesis of *R. solani*, the dsRNA treated mycelia were inoculated in 30 days old tomato plants and incubated in the infection chamber at 28°C (Ghosh et al. 2021). The disease symptom was recorded at 3 dpi and disease index as well as pathogen load was calculated, as described previously^21^. Atleast five plants (three leaves per plant) were analysed for each treatment and the experiment was independently repeated three times (biological replicates).

## Data availability

The genome assembly has been submitted to NCBI with bioproject ID PRJNA715598.

## Acknowledgements

AF and KT acknowledge fellowship from CSIR, Govt of India and DBT, Govt. of India, respectively. DC acknowledges JC bose fellowship (JCB/2020/000014) from Science and Engineering Research Board (SERB), Govt of India. GJ acknowledges core research grant from National Institute of Plant Genome Research, India, Swarna Jayanti Fellowship (SB/SJF/2020-21/01) from SERB, Govt of India and NIPGR flagship program (102/IFD/SAN/763/2019-20; supported by DBT). The funders had no role in study design, data collection and analysis, decision to publish, or preparation of the manuscript. The authors are also thankful to DBT-eLibrary Consortium (DeLCON) for providing access to e-resources.

## Author contributions

GJ has conceived the work, coordinated the entire process and has supervised wet-lab experiments. DC has planned and supervised the computational part of the study. AF has performed the genome assembly and most of the computational analysis. SG has characterized *R. solani* strains, prepared samples for genome sequencing, and identified genes related to pathogenicity. KT and SG have standardized dsRNA based silencing of *R. solani* genes. KT and MR have carried out functional characterization and expression analysis of *R. solani* genes. AF, KT and MR have assembled the figures and tables. VP, RMS, CP, GSL, CK and MSP have contributed 32 *R. solani* strains, generated their passport data and provided comments on the mansucript. NS has developed *R. solani* genome database and carried out phylogenetic analysis. SG, DC and GJ have written the manuscript and all authors have approved it.

