## supplementary information for "Genome duplication and transposon mediated gene alteration shapes the pathogenicity of *Rhizoctonia solani* AG1-IA"

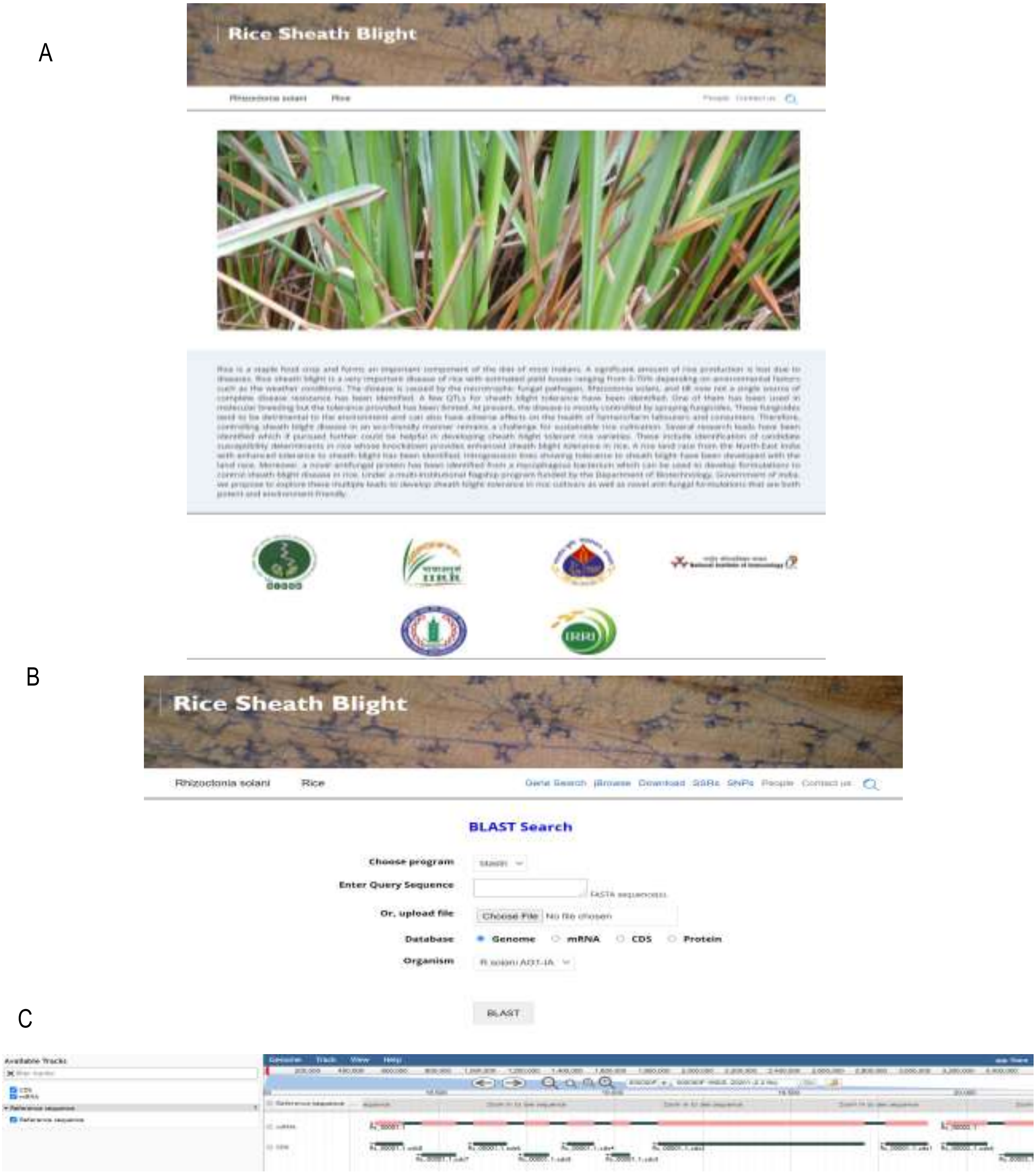


Fig. S1. **Rice sheath blight (RSB) database.** A. Home page; B. Blast search interface; C. Jbrowser for genome visualization

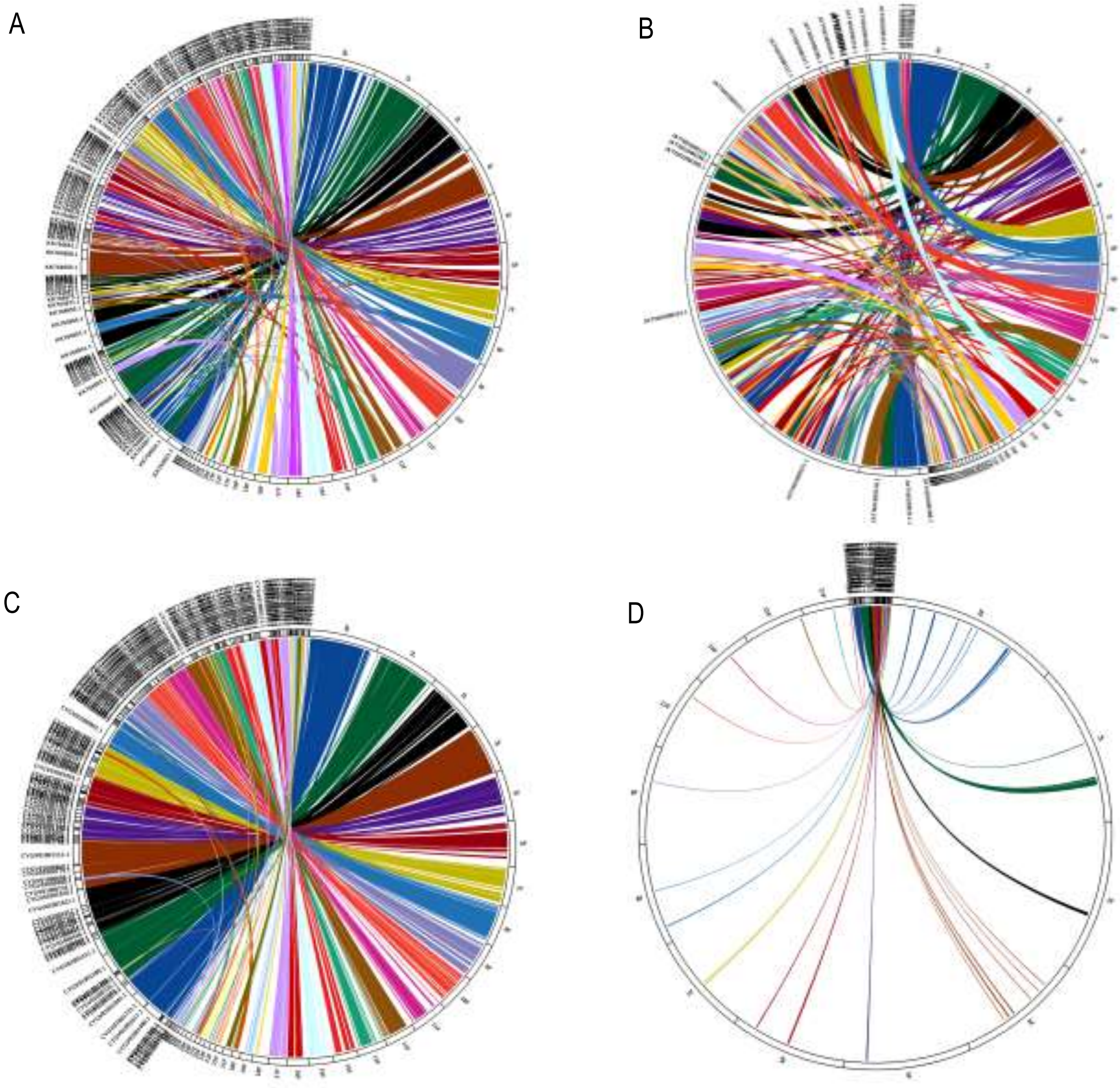


Fig. S2. Synteny between AG1IA and A. AG8;B. AG3; C. AG22IIIB; D. AG1IB

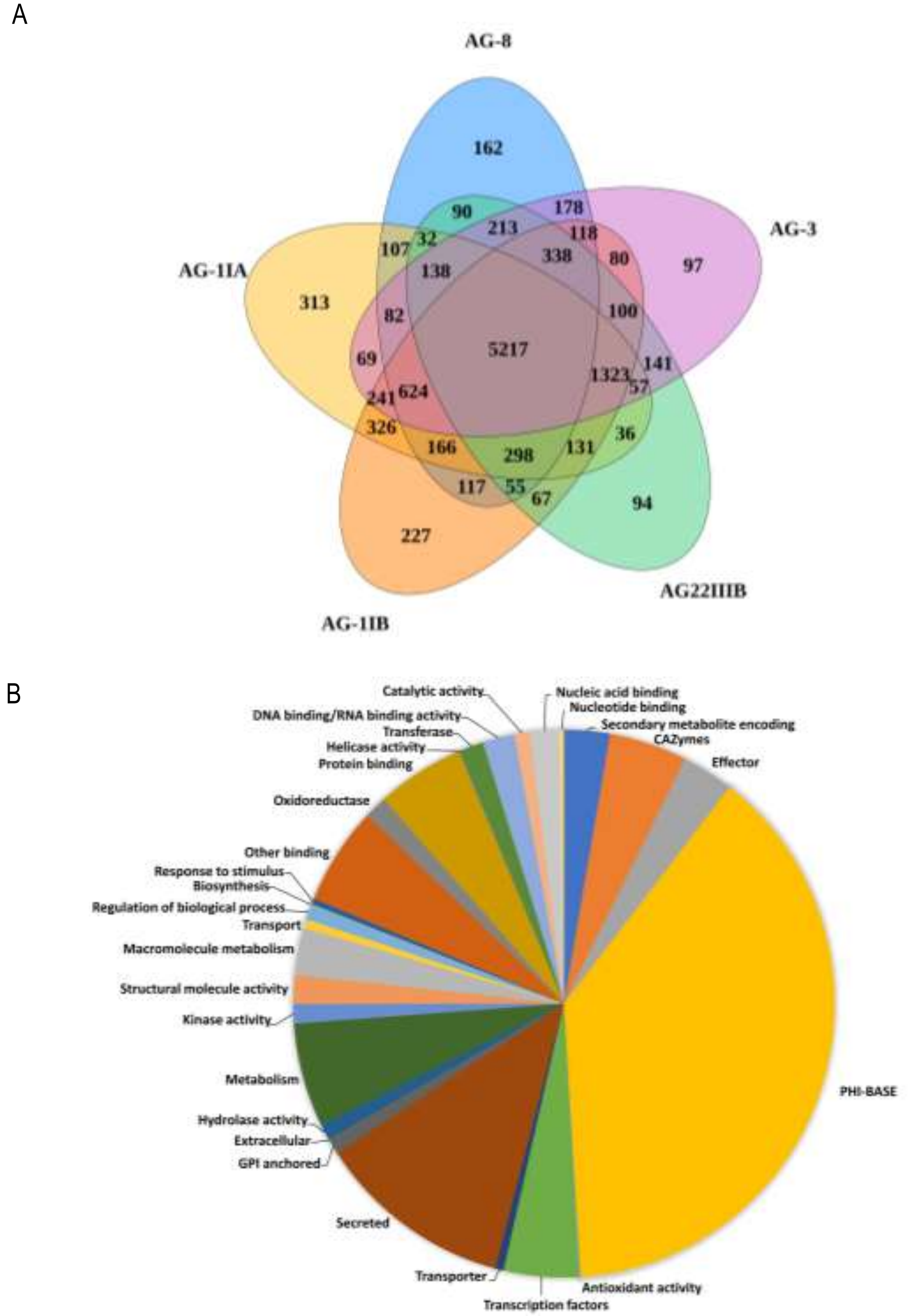


Fig. S3. A. Venn diagram showing distribution of gene families among the genome assemblies of *R. solani* strains. Comparative analysis revealed that a total of 73015 genes from five strains are clustered into 20840 families. The number of gene families that are unique and are shared among different strains are indicated, B. The Pi chart shows distribution of functions of protein-coding genes encoded by the *R. solani* AG1-IA genome assembly.

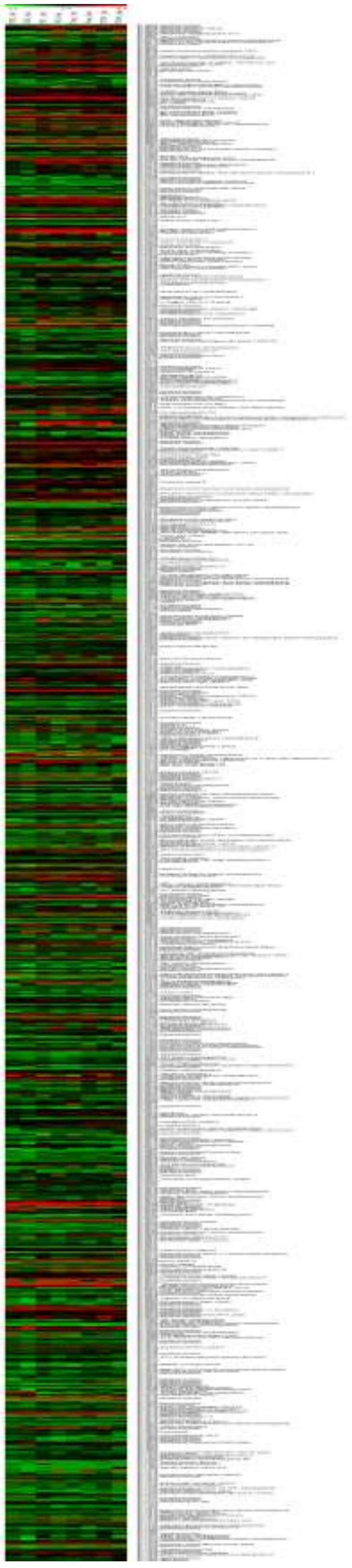

Fig. S4. RNA-seq based expression analysis of *R. solani* AG1IA strain BRS1 specific genes

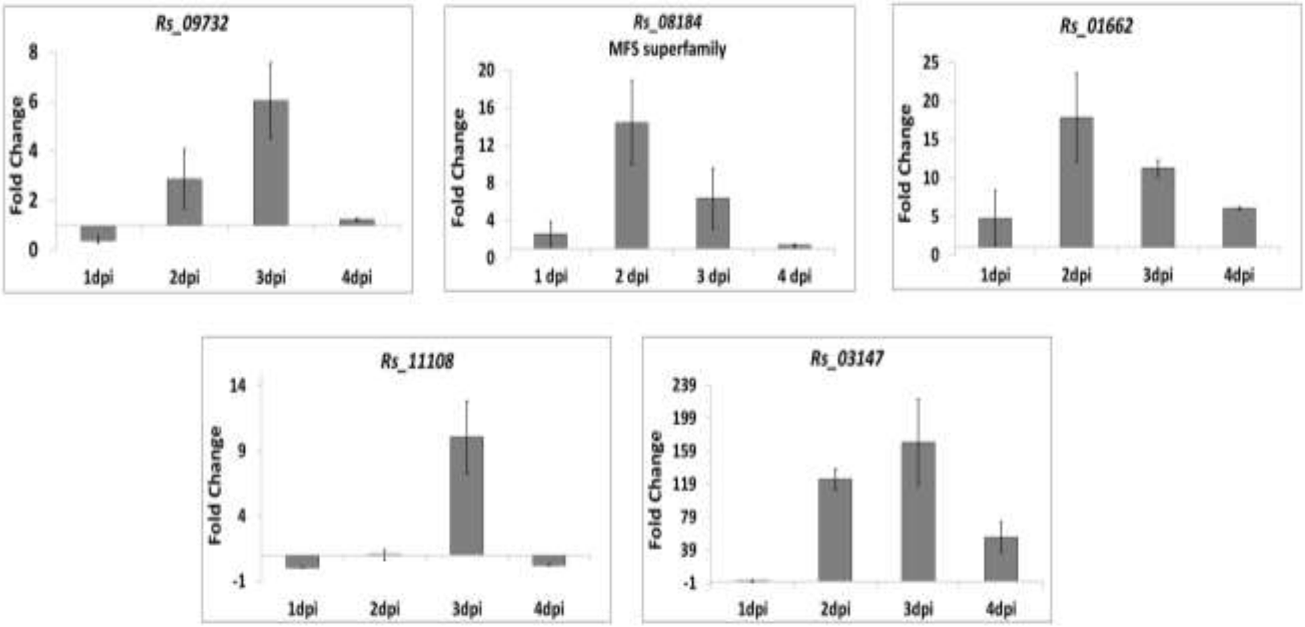


Fig. S5. Expression analysis of *R. solani* AG1-IA strain BRS1 unique genes during pathogenesis in rice (cv. PB1). The gene expression at indicated time points was quantified with respect to 0 dpi samples. The predicted molecular function/ domain are indicated for each gene. *R. solani 18S rRNA* gene was used as endogenous control. Data represents mean value of three biological replicates and error bars indicates standard error of the mean. MFS= Major Facilitator Superfamily.

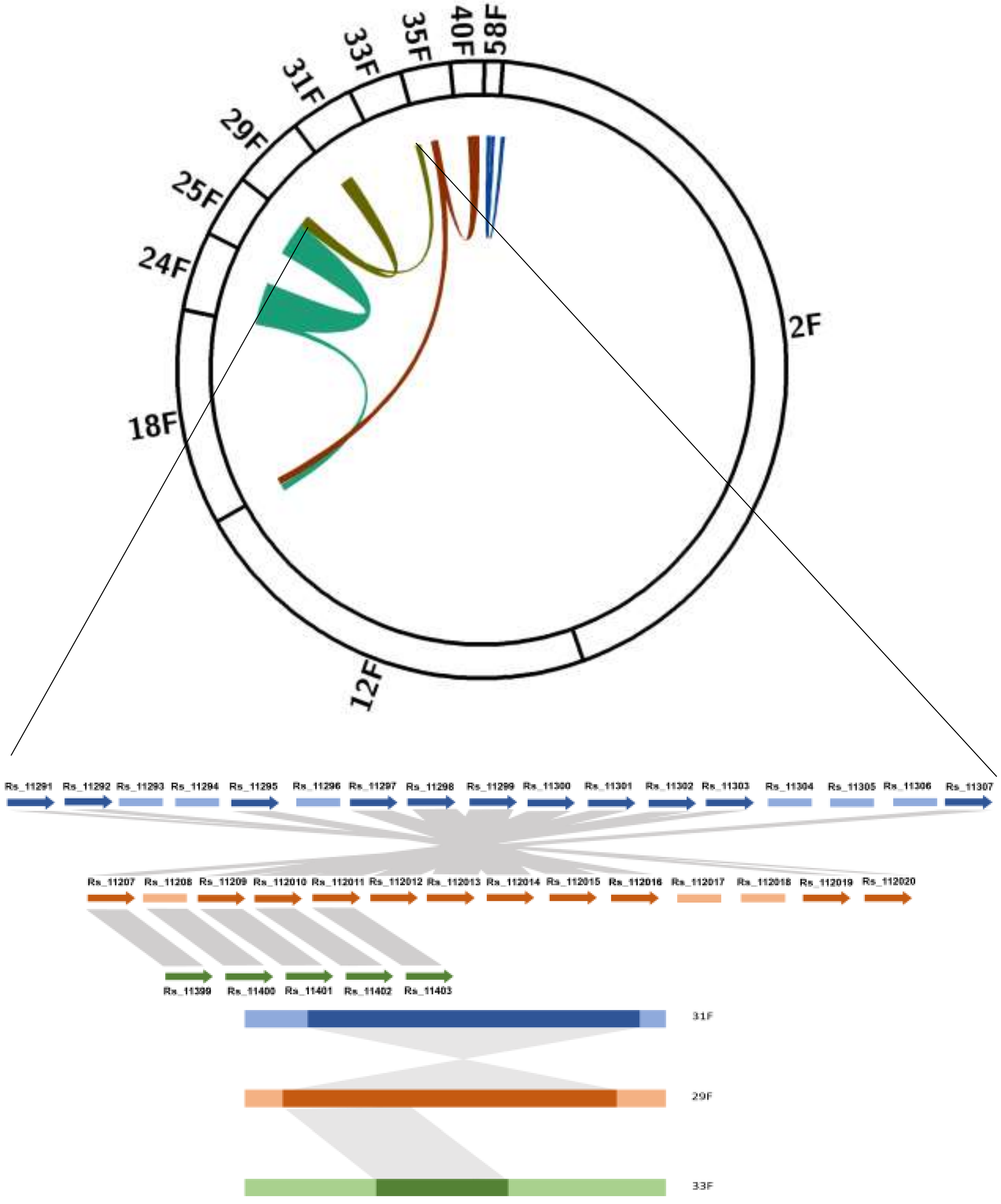


Fig. S6. Depiction of four triplicated paralogous blocks of AG1IA genome

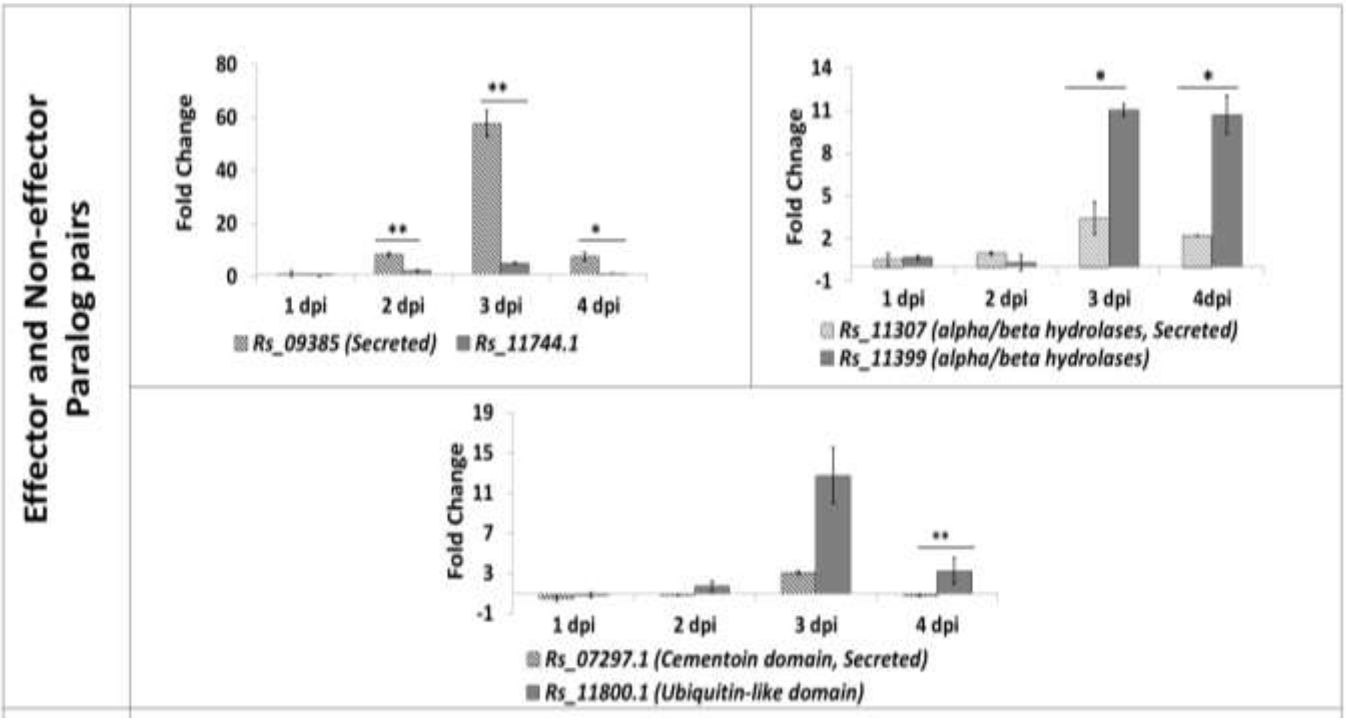


Fig. S7. Expression analysis of *R. solani* AG1-IA paralog pairs during pathogenesis in rice (cv. PB1). The expression at indicated time points was quantified with respect to 0 dpi samples. The paralog pairs were categorized in three categories including Effector or non-effector (presence or absence of predicted secretory signal) and their importance in *R. solani* selection. Data represents mean value of three biological replicates and error bars indicates standard error of the mean.

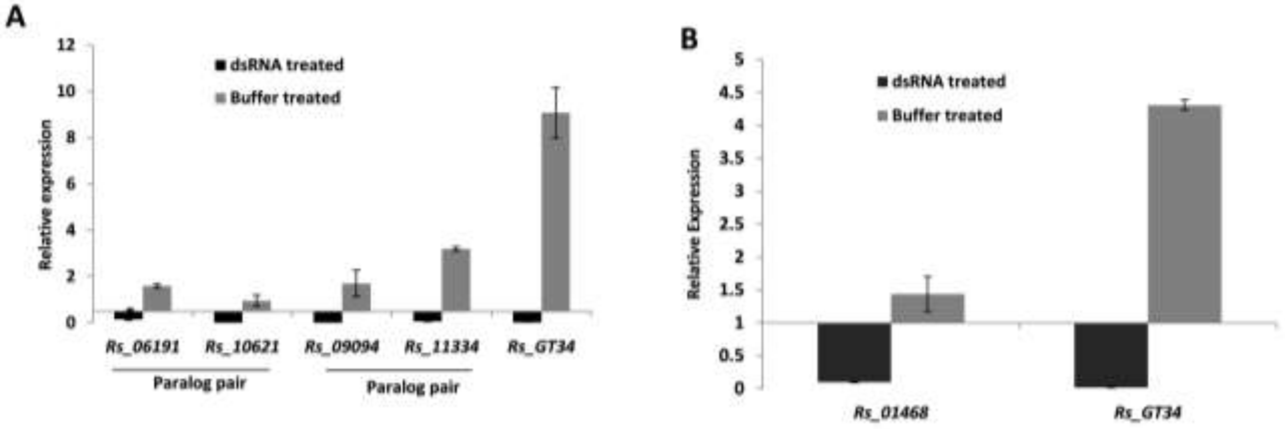


Fig. S8. dsRNA mediated silencing of *R. solani* genes. qRT-PCR based expression analysis of A. Paralog pairs with domain variation and B. *Rs_01468* (gene under positive selection) gene of *R. solani* upon silencing compared to control, at 3 dpi. *18S rRNA* of *R. solani* was used as endogenous control. Data represents mean values ± standard error of three biological replicates.

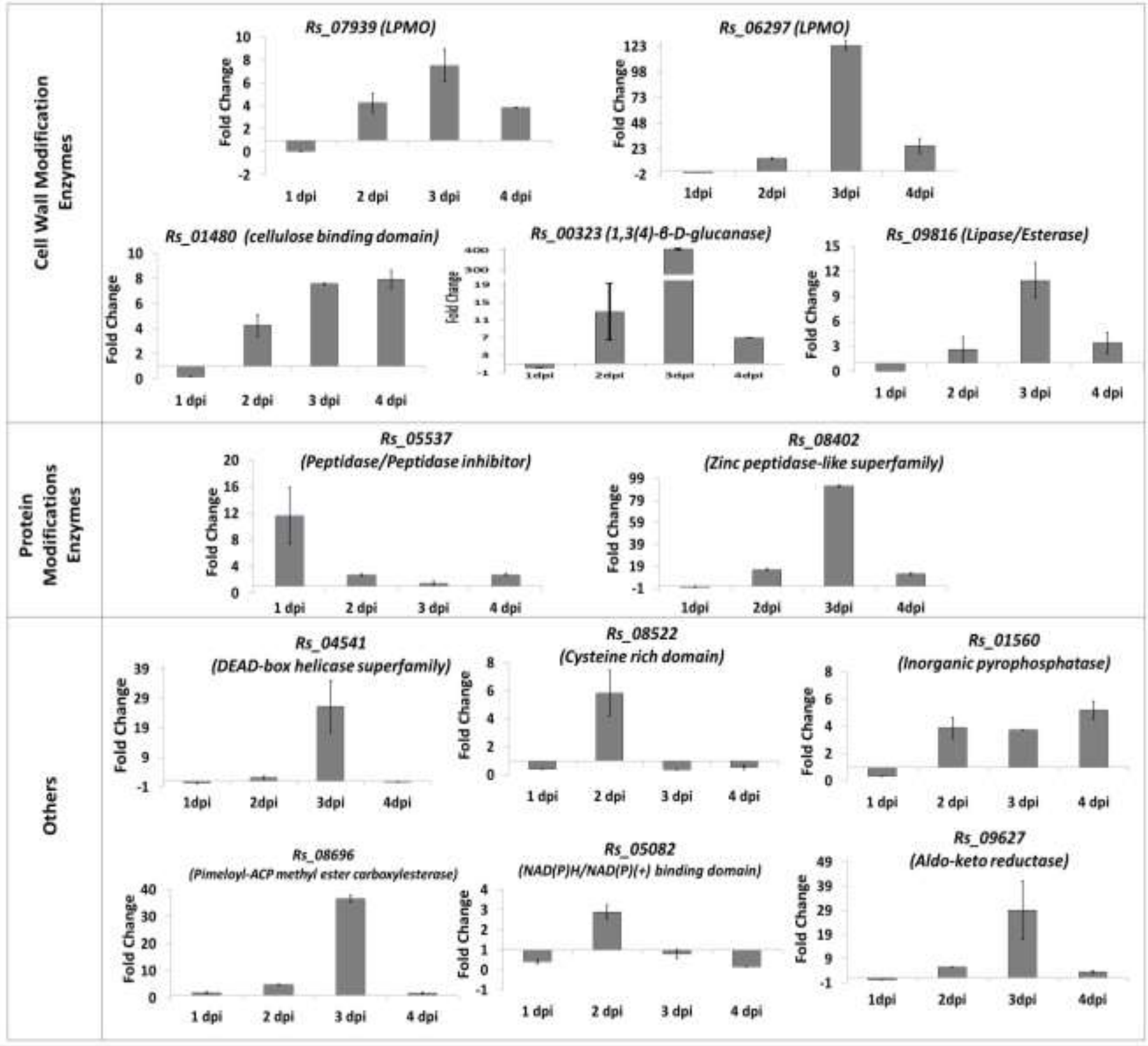


Fig. S9. **Expression analysis of *R. solani* AG1-IA genes under positive selection during pathogenesis in rice (cv. PB1).** The expression at indicated time points was quantified with respect to 0 dpi samples. The predicted molecular function/ domain for each gene is indicated. *18S rRNA of R. solani* was used as endogenous control. Data represents mean value of three biological replicates and error bars indicates standard error of the mean. LPMO=Lytic polysaccaharide monoxygenase, ACP=Acyl carrier protein

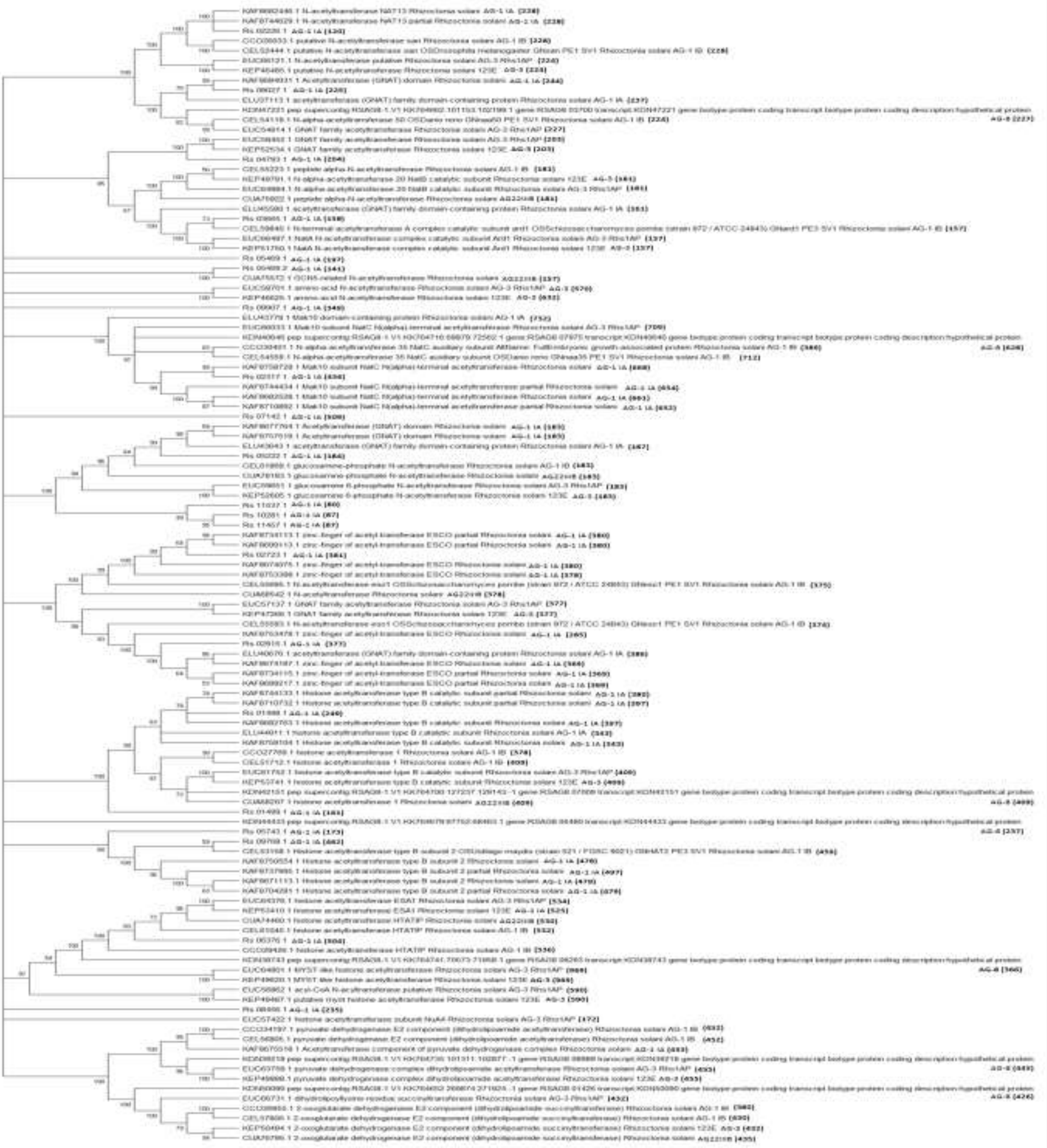

Fig. S10. **Phylogenetic analysis of *R. solani* GNAT gene family**. Unrooted phylogenetic tree of GNAT proteins constructed by neighbor joining method with 1000 bootstraps using MEGA X tool. Numbers on branches represent bootstrap values (based on 1000 replications), only the values ≥50% are presented.
